# Effects of microplastics on Daphnia-associated microbiomes in situ and in vitro

**DOI:** 10.1101/2024.06.28.601137

**Authors:** Anna Krzynowek, Broos Van de Moortel, Nikola Pichler, Isabel Vanoverberghe, Johanna Lapere, Liliana M. Jenisch, Daphné Deloof, Wim Thielemans, Koenraad Muylaert, Michiel Dusselier, Dirk Springael, Karoline Faust, Ellen Decaestecker

## Abstract

Microplastics (MP) pollution in aquatic environments is a growing global concern. MP, defined as plastic fragments smaller than 5mm, accumulate in freshwater reservoirs, especially those located in urban areas, impacting the resident biota. This study investigated the effects of MP on the performance and microbiome of Daphnia, a keystone organism in freshwater ecosystems, through both in situ sampling of freshwater ponds and a controlled 23-day in vitro exposure experiment. Using 16S rRNA gene sequencing and whole-genome shotgun sequencing, the microbiome community composition and functional capacity was analysed and correlated with MP pollution levels. Urban ponds showed higher MP concentrations in both water and sediment than natural ponds with significant differences in MP composition. Bacterioplankton communities were more diverse and richer than the Daphnia-associated microbiomes. Overall, the in situ study showed that the composition of the Daphnia-associated community co-varied with high MP levels but also with temperature and redox potential. Moreover, the functional analysis showed increased relative abundances of PET degradation enzymes and antibiotic resistance genes (ARGs) in microbiomes from high-MP ponds. In the in vitro experiment, the bacterioplankton inoculum source significantly influenced Daphnia survival and microbiome composition. Daphnia exposed to high MP concentrations inoculated with bacterioplankton pre-exposed to MP exhibited significantly higher survival rates, suggesting potential adaptive benefits from MP-associated microbiomes. Network analysis identified specific taxa associated with MP within the Daphnia microbiome. Our study suggests adaptive responses of freshwater host-associated microbiomes to MP pollution including biodegradation with potential benefits for the host.

## INTRODUCTION

By 2030, the world’s aquatic environment is predicted to accommodate more than 80 metric tons of plastic waste (Borrelle *et al*., 2020; Tian *et al*., 2022). **Microplastics (MP)**, i.e., plastic particles smaller than 5 mm originating from either fragmentation of larger plastic materials or as such released in the environment, have become of major environmental concern not only in marine but also in freshwater ecosystems (Raju *et al*., 2023). In particular, stagnant ponds near densely populated areas are prone to high plastic deposition burdens showing concentrations of up to 19,860 particles/m^3^ (Nava *et al*., 2023; Xiong *et al*., 2018; Raju *et al*., 2023). While studies have mainly focused on the impact of MP on marine organisms, MP also affect **freshwater** organisms at different levels of the trophic chain (Yuan et al., 2019). This includes members of the zooplankton genera ***Daphnia*** which occupy an important lower trophic level within aquatic food webs. *Daphnia* feeds on small suspended particles in the water and sediments in freshwater ecosystems resulting into MP accumulation upon ingestion (Elizalde-Velázquez *et al*., 2020; Jemec *et al*., 2016; Gökçe *et al*., 2022). Acute effects of MP exposure have been observed in *Daphnia,* and include molecular responses (Trotter *et al*., 2021; Liu *et al*., 2020; Ma *et al*., 2016), effects on survival (Liu *et al*., 2020; Jaikumar *et al*., 2018; Cui *et al*., 2017), reproduction (Liu *et al*., 2020; Cui *et al*., 2017; Aljaibachi *et al*., 2020; Zimmermann *et al*., 2020; De Felice *et al*., 2019), feeding rates (Jemec *et al*., 2016; Rist *et al*., 2017), body mass (Cui *et al*., 2017) and mobility (Jemec *et al*., 2016; Rehse S *et al*., 2016). *Daphnia*’s carapax, body cavities, gut, and feeding apparatus are colonised by bacteria, collectively defined as **the microbiome** (Qi *et al*., 2009; Eckert *et al*., 2014; Cooper *et al*., 2020). The assembly of this microbiome involves complex interactions with the surrounding bacterioplankton and is influenced by the host’s diet (Li *et al*., 2021; Callens *et al*., 2016), genetic background (Frankel *et al*., 2020; Macke *et al*., 2017; Callens *et al*. 2020), and environmental conditions (Li *et al*., 2019; Sullam *et al*., 2018). The host microbiome contributes to the host’s fitness (Sison-Mangus *et al*. 2015). Moreover, core members of *Daphnia*’s microbiome have been identified (Cooper *et al*., 2020), with certain taxa positively correlated with increased host body size, growth, and survival (Peerakietkhajorn *et al*., 2015, 2016; Sison-Mangus *et al*., 2015). Additionally, the gut microbiome is suggested to play a role in the degradation of toxins, such as those produced by cyanobacteria (Macke *et al*., 2017; Houwenhuyse *et al*., 2021). Thus, *Daphnia*’s microbiome not only contributes to host survival but may enhance the host’s ability to adapt to and restrain environmental stressors.

Studies that address the impact of MP pollution on the *Daphnia* microbiome are limited. A recent short-term study on polystyrene (PS) particles showed a transfer of microbiota from *Daphnia magna* to MP biofilms (Gorokhova *et al*., 2021), which are essential hubs for plastic polymer biodegradation (Howard *et al*., 2023; Adam *et al*., 2024). *Vice versa*, the biofilm community may also interact with the already established microbiome community. In this scenario, the so-called **plastisphere** community may lead to the transfer of specific members to the particle-feeding zooplankton microbiomes. With the transfer of these members comes the transfer of some genes associated with microplastics (MP). These include plastic polymer degradation enzymes and antibiotic resistance genes (ARGs), which have been demonstrated to spread via MP-associated biofilms (Kaur *et al*., 2022; Zhao *et al*., 2024).

Here, we studied the effects of MP on *Daphnia’s* microbiome *in situ* by sampling freshwater reservoirs and *in vitro* by conducting a 23-day MP-exposure experiment (**Figure 1**). In the *in situ* study, we measured the prevalence of MP across urban and natural freshwater ponds in Flanders, Belgium and used 16S rRNA marker gene and whole genome shotgun sequencing to elucidate microbial community composition and functional capacity of the bacterioplankton and native *Daphnia* populations, which were then correlated with MP pollution levels. The field campaign results were compared to those of the *in vitro* exposure experiment in which we cultured different *Daphnia magna* clones with PET, PLA (polyester), and Nylon (polyamide) MP for 23 days while monitoring *Daphnia* survival. The microbiome community composition was resolved using 16S rRNA marker gene sequencing and associations of taxa with MP exposure were investigated.

**Figure 1.**
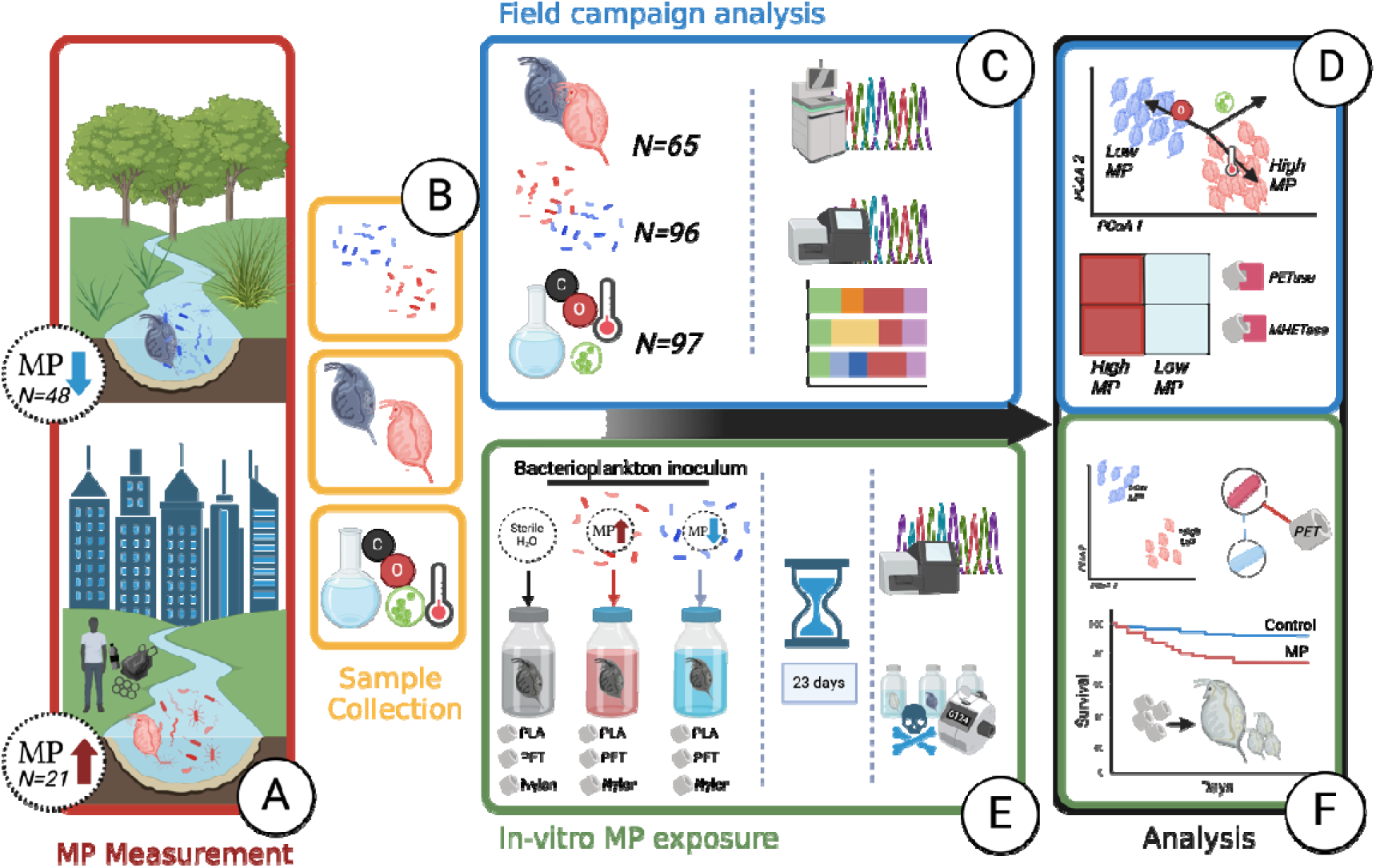
Graphical overview of the study design and experiments. **A.** Samples of sediment cores and water were collected from two types of ponds, natural lakes (N=24,24 for water, sediment) and urban ponds (N=12,9 for water, sediment), for MP quantification. Additionally, the following data were collected: **B.** Pond bacterioplankton, local *Daphniidae* populations, and environmental factors. **C.** Water parameters (N=97) were compared across sampled ponds; bacterioplankton (N=96) and *Daphniidae* microbiome samples (N=65), were sequenced with 16S rRNA for microbial community analysis and whole genome shotgun sequencing at high depth (1 million reads per sample) for functional analysis. **D.** The community composition of bacterioplankton and whole-Daphnia microbiomes was compared between high and low MP ponds, while accounting for sampled environmental factors. Furthermore, the functional capacity of host microbiomes, including the average abundance of PET-degrading enzymes and Antibiotic resistance genes (ARGs) was compared between the two environments. **E.** In the short-term exposure experiment, three Daphnia magna genotype clones were exposed to PET, PLA, and Nylon microplastic fibres for 23 days. The exposure jars were inoculated with bacterioplankton from either the high-MP Blauwe Poort pond or the low-MP De Gavers pond, collected during the field campaign. After 23 days, the DNA from both animals and bacterioplankton was extracted and sequenced with 16S rRNA marker gene. Data on population and animal survival were collected throughout the experiment. **F.** The community composition of bacterioplankton and whole-Daphnia microbiomes was compared between different MP exposures (PET, PLA, Nylon) and non-exposed animals (Negative Control). Additionally, an association network between taxa and MP was constructed to identify taxa significantly associated with specific MP. Finally, *Daphnia* survival in response to MP exposure was analysed throughout the experiment.

## MATERIALS AND METHODS

### 1. Field sampling campaign

#### 1.1. Pond sampling

Bacterioplankton, *Daphnia* population and MP were sampled from twelve ponds located in Flanders, Belgium (**Supplementary Figure 1**). The ponds were categorised as natural lakes or city ponds based on a location (**Supplementary table 2**). On site measurements of temperature, pH, oxygen, redox, and conductivity were carried out in triplicate using a Hach HQ40d multi-meter. NPOC concentrations were measured in the lab with a Shimadzu TOC-L analyzer (**Supplementary table 3**). A total of 161 samples of *Daphnia* (65) and bacterioplankton (96) were collected. To recover *Daphnia*, each 30L of sampled water was filtered through a 300µm zooplankton net, rinsed, dried, and frozen at -80°C. For bacterioplankton, 250mL water samples were collected at three depths (bottom, middle, top) and filtered through 0.22μm sterile filters which were subsequently frozen at -80°C until processing. MP were sampled from both sediment and water columns in triplicate and within 1-metre distance from each other. For water, 150L first sieved through stacked metal sieves (1000µm, 500µm, 250µm, 64µm) and MP (64µm-250µm) were collected in a glass jar. Sediment samples from the top 5 cm of solid sediment (where possible) or around 500mL of loose sediment. All sediment and water samples were handled using non-plastic equipment.

#### 1.2. MP quantification and characterization

MP particles from the water in the glass jars and from sediment samples were recovered as described by Meyers *et al*., (2024a, 2024c). The process involved three steps: digestion of organic matter with H2O2, staining with Nile Red-dye, and applying a machine-learning algorithm to distinguish and count plastic particles (**Supplementary materials and methods).**

#### 1.3. Molecular biology

DNA from bacterioplankton and whole *Daphnia* samples was extracted with the Powersoil Pro (Qiagen) and NucleoSpin (Macherey-Nagel) kits, respectively, following the manufacturers’ protocols. Extracted DNA was resuspended in a 20µL elution buffer, quantified with a NanoPhotometer N60 and used for 16S rRNA gene amplicon and whole-genome shotgun sequencing. The V4 region of the 16S rRNA gene was amplified using a nested PCR strategy as detailed in **Supplementary materials and methods**. Shotgun libraries were prepared with the PCR-free Kapa Hyperprep kit and sequenced on the Illumina NovaSeq 6000 platform at 100Gb per sample at the KULeuven Genomics Core, Belgium.

#### 1.4. Analysis of MP distribution in nature

Average MP concentrations between urban and natural pond categories were compared in sediment and water samples. All calculations were performed in R-studio version 4.3.1. Bar charts were generated using ggplot2 and ggpubr with error bars representing the standard error of the mean. The data was assessed for normality using the Shapiro-Wilk test followed by a two-sample t-test to check for statistically significant differences between average MP in urban and natural ponds. The differences in composition were tested with Multivariate Analysis of Variance (MANOVA). The analysis was conducted using Python’s (ver. 3.11.7) statsmodels library, with MANOVA providing four test statistics: Wilks’ lambda, Pillai’s trace, Hotelling-Lawley trace, and Roy’s greatest root, to assess the multivariate effect of pond category on the composition of MP concentrations.

#### 1.5. Analysis of 16S rRNA amplicon sequencing for quantifying differences in community composition of Daphnia and bacterioplankton samples

The quality of demultiplexed paired-end reads was assessed using FastQC (Andrews, 2010). The run generated 10,832,180 raw sequences (min 7781, max 55,923 per sample). Paired-end sequences were filtered, denoised, and dereplicated with DADA2 (Callahan et al., 2016) in R-studio. Taxonomy was assigned using the Silva 138.1 SSU Ref NR 99 database (Quast et al. 2013). Reads labelled as “mitochondria” or “chloroplast” were removed. The ASV table, taxonomy, and metadata were imported into a phyloseq object (v. 1.44.0, McMurdie and Holmes 2013). A rarefaction curve was constructed using the vegan package (v. 2.6–4, Oksanen et al. 2013), and samples were rarefied to the depth of the sample with the fewest reads (4,664), removing 2007 ASVs. Bray-Curtis dissimilarities were computed for ordination. Environmental variable correlations to the axis were computed with the envfit function (vegan package), plotting vectors with p-value < 0.05. For distance-based redundancy analysis (db-RDA), highly collinear variables (correlation >0.7) were removed and remaining ones standardised. The significance of the association between environmental variables and community composition was tested with permutation tests (alpha=0.05). Diversity indices were calculated using alpha and boxplot_alpha functions from the microbiome package (v. 1.22.0, Leo Lahti, Sudarshan Shetty et al. 2017).

#### 1.6. Analysis of whole-genome shotgun of whole Daphnia-microbiome samples

Demultiplexed paired-end reads were quality-checked using FastQC, adapter sequences were trimmed, and reads with a Phred score below 30 were filtered out using Fastp (Chen et al., 2018), resulting in 12.59 billion high-quality reads (14,330,348 to 261,293,254 per sample). Eukaryotic sequences were identified and removed with Kraken2 v.2.1.3 (Wood et al., 2019) using the NCBI non-redundant nucleotide database. Reads mapping to genomes of *Cladocera*, *Daphnia*, microsporidia, humans, and freshwater algae were excluded using Bowtie2 (Langmead & Salzberg, 2012), and unmapped pairs were extracted with Samtools (Li et al., 2009). Bacterial reads were co-assembled using MEGAHIT (Li et al., 2015). Contigs were checked against the Refseq bacteria database with Kraken2 and only bacterial contigs were analysed. Open Reading Frames (ORFs) were predicted with Prodigal (Hyatt et al., 2010). Quality-filtered reads were aligned to contigs using Bowtie2 and alignment statistics were obtained with Samtools. FeatureCounts (Liao et al., 2014) extracted read counts for each ORF. Gene abundances were normalised using the adapted RPKM formula (Dillies et al., 2012):

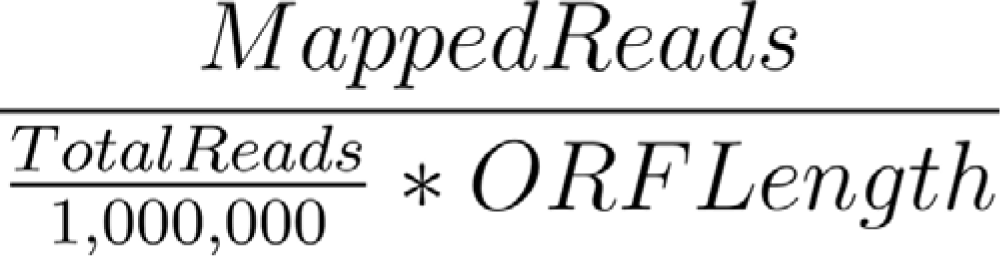

Proteins were annotated with eggNOG-mapper v2.1.6 (Huerta-Cepas et al., 2019), and abundances were derived using FeatureCounts. Differences in gene distributions were visualised with PCoA ordination on the Bray-Curtis dissimilarity matrix. Functional enrichment between high and low MP groups was assessed using DESeq2 (Love et al., 2014). PET degradation pathways were compared by mapping KO identifiers against the KEGG database (Kanehisa et al., 2015). ARGs were identified by aligning protein sequences against the CARD database (Jia et al., 2017) using BLASTP (Altschul et al., 1990) with stringent criteria.

### 2. In vitro Daphnia magna exposures to MP

#### 2.1. MP fibre generation

Three types of polymers were used, i.e, the poorly biodegradable polyethylene terephthalate (PET) and polyamide 6,6 (Nylon) and the (under industrial composting conditions) biodegradable alternative polylactic acid (PLA) (Narmon et al., 2022). All three polymers are used in textile manufacturing and their MP are found in waste waters from textile fabrics and household washing activities. PET and Nylon fibres (diameter 0,01 mm) were procured from Goodfellow (GF01552395 and GF15603331, respectively) and PLA filaments (0.05 mm diameter) were kindly provided by Sioen Industries (Ardooie, Belgium). Uniformly sized microfiber preparations were obtained as described by Cole et al. (2016), including solidifying them at -80°C, cutting into 50µm pieces, and processing with heating and vacuum filtration (**Supplementary materials and methods**). The fibres were sterilised with ethanol and stored in a sterile container until use.

#### 2.2. Determining optimal MP concentrations

To determine an optimal concentration of MP in the exposure experiment, concentrations of 12 mg/L, 50 mg/L, and 100 mg/L of PET, PLA and Nylon fibres were tested for acute (48 hours) and chronic (23 days) effects on three *Daphnia magna* clones (KNO15.04, F, and OM2.11) (**Supplementary materials and methods**). A concentration of 2.5±0.1 mg/L was selected for the main experiment which was significantly above typical field conditions (0.021 mg/L for Flemish surface water) (Vercauteren et al. 2021).

#### 2.3. Daphnia magna clones

For the exposure experiment, three *Daphnia magna* clonal lineages were used: F (Barata et al., 2017), KNO15.04 (from a fishless farm pond near Knokke, 51°20′05.62″ N, 03°20′53.63″ E), and BP (collected from the “Blauwe Poort” pond). These genotypes were maintained for many generations at the IRF life sciences laboratory in Kortrijk as reported (Coone *et al*., 2023). At the start of the experiment, 10 hatched neonate *Daphnia* individuals of less than 24 hours old were transferred with glass pipettes from the Petri dishes into the experimental jars.

#### 2.4. Experimental design

To compare the effects of bacterioplankton inocula from low MP and high MP ponds on the *Daphnia* microbiome community and on *Daphnia* survival we used pond water taken as 18 L samples from De Gavers (DG) lake and from the plastic-polluted city pond Blauwe Poort (BP) as bacterioplankton inocula in the exposure experiments. Temperature (°C), dissolved oxygen (mg/L), liquid dissolved oxygen (%), pH, conductivity (µs/cm) and redox (mV) were determined in the two waters before the experiment (**Supplementary materials and methods**). The experiment was set up in 500mL glass jars in triplicate for genotype (F, KNO 15.04, BH, No *Daphnia*) x MP (PET, Nylon, PLA, No MP) x pond microbial inoculum (De Gavers, Blauwe Poort, Sterilised H2O) yielding 99 jars in total (**Supplementary materials and methods**). After sterilising the jars with ethanol and washing them with purified MQ water, they were filled with 400 mL of the bacterioplankton inoculum and 10 *Daphnia* individuals as well as 1±0,1 mg of microfibers were added. All jars were covered with tin foil to prevent water evaporation and contamination with external microfibers.

#### 2.5. Data collection

The survival of 10 *Daphnia* individuals was checked every two days. Body size was measured on days 7 and 14 for five randomly selected adults using a stereomicroscope (Olympus SZX10) and ImageJ software. The date and size of the first brood were recorded by measuring five random juveniles. On days 3, 7, and 14, five *Daphnia* were examined for ingested MP fibres using a light microscope (Olympus BX51) **(Supplementary materials and methods)**. The experiment lasted 23 days, with *Daphnia* fed at a feeding regime of 2 x 10^5^ cells/ml of *Chlorella vulgaris* every two days and of 1 x 10^5^ cells/ml towards the end. Jars were shaken every two days. On day 23, five adults and five juveniles from each jar were harvested, dried, and stored at -80 °C for microbiome analysis. For bacterioplankton analysis, 50 mL of water was filtered through a sterile 0.22 μm filter and the recovered cells stored at -80 °C.

#### 2.6. Molecular biology

A total of 246 samples were processed for DNA extraction. The samples were split into *Bacterioplankton* (101 samples) and *Daphnia* (Juveniles: 41 samples, Adults: 102 samples). Protocols for DNA extraction and 16S rRNA gene sequencing were identical to those used for the samples collected from the field.

#### 2.7. 16S rRNA gene metabarcoding of the MP exposure experiment

For 16S rRNA gene metabarcoding, demultiplexed paired-end reads were imported and assessed with FastQC, resulting in 6,901,562 raw sequences (min 92, max 23,577 per sample). Sequences were filtered, denoised, and dereplicated with DADA2 (Callahan et al., 2019) in R-studio. Taxonomy was assigned using the Silva 138.1 SSU Ref NR 99 database (Quast et al. 2013). Contaminating taxa were removed using the microDecon package (v. 1.0.2, McKnight et al. 2019). Two extraction controls were used to detect contamination. Sequences annotated as “mitochondria” or “chloroplast” were removed. The ASV table, taxonomy, and metadata were imported into R with the phyloseq package (v1.44.0, McMurdie&Holmes, 2013). Rarefaction curves were constructed, and ASV counts were rarefied to 2,494 reads, resulting in the loss of 3,010 ASVs. Significant effects were assessed using permutational multivariate ANOVA (permanova) with homogeneity of variances evaluated using functions from the vegan package (Okasen et al., 2013).

#### 2.8. Survival analysis of Daphnia magna in response to MP exposure

Survival curves were calculated based on four time points with the Kaplan-Meier estimator, using the survfit() function in the survival R package. Survival curves were stratified by pond type, fibre type, and *Daphnia magna* genotype using ggsurvplot(), with significance tested by a log-rank test (alpha = 0.05). A Cox proportional hazards model (coxph()) assessed multiple factors’ impact on survival, including pond type, genotype, fibre type, and replicate. Statistical significance was evaluated with Wald tests, likelihood ratio tests, and score tests. The impact of pond type, genotype, and fibre type on *Daphnia* body size was assessed with generalised linear models (GLMs) implemented in the glm package, using a Gamma distribution and log link for body size and a Poisson distribution for the age at first brood. Models’ goodness of fit was assessed by comparing null and residual deviances, Akaike Information Criterion (AIC), and Fisher Scoring iterations.

#### 2.9. Network analysis and differential taxa enrichment analysis

Before network inference, the rarefied ASV counts were first subjected to a prevalence filter of 10% to remove sparse taxa using custom functions in R. Categorical metadata variables were one-hot encoded. The network for *Daphnia* exposure communities were built using FlashWeave version 0.19.2 (Tackmann *et al*., 2019). Both networks were built in sensitive mode. The networks were exported in gml format and visualised with Cytoscape v3.10.1. All p-values were FDR-corrected using the Benjamini-Hochberg procedure (Benjamini & Hochberg, 1995).

## RESULTS

### Urban ponds accumulate more MP than natural lakes

Higher MP concentrations were found in city pond sediments compared to natural pond sediments (Two sample T-test, p-value < 0.01) (**Figure 2A**). At the level of individual ponds, two urban ponds, *Kluizen* and *Blauwe Poort* (BP), showed the highest MP concentrations in the sediment with an average of 21.56 and 10.31 MP per gram of sediment, respectively. *Meer van Rotselaar* and *Reserve Kulak*, two natural lakes, contained the lowest concentrations with an average of 0.08 and 0.16 MP per gram of sediment, respectively (**Supplementary file 8**). Similarly, in the water samples, higher MP concentrations were found in the urban ponds than in the natural lakes (Wilcoxon rank sum test, p-value < 0.01) **(Figure 2C**) with the highest average MP concentration measured in an artificial city pond (*Citadelpark,* 1366 MP in mL of sample) and the lowest in a natural lake (*Meer van Rotselaa*r, 22 MP per mL of water) (**Supplementary file 8**). The composition of MP varied across the samples per pond (water vs sediment) and pond type (urban vs natural). Polyethylene/Polypropylene (PE/PP) were the most abundant MP types in the urban pond sediments followed by Polystyrene (PS) and PET/Polyester (**Figure 2B**). In contrast, PET/Polyester was more abundant in water samples (**2B,D**).

**Figure 2.**
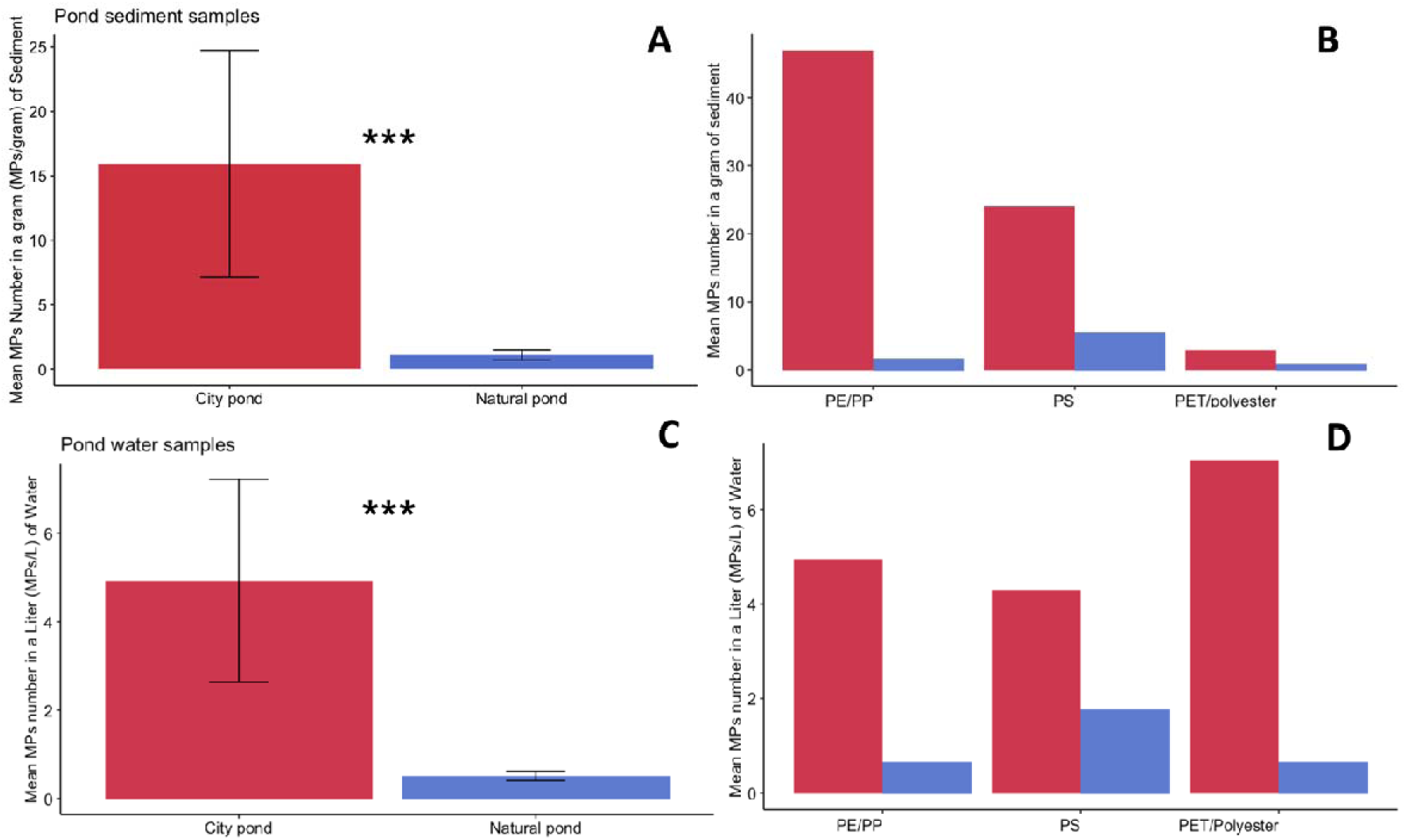
MP analysis collected from the pond water and sediment. **A.** Difference between the average concentration of MP measured in the sediment from urban ponds (Red) and natural lakes (Blue) with standard error of the mean plotted on the bars. **B.** Average concentrations of MP per gram of sediment (n=3) for each MP type measured in the study. **C.** Difference between the average concentration of MP measured in the water samples from artificial city ponds (Red) and natural lakes (Blue) with standard error of the mean plotted on the bars. **D.** Average concentrations of MP per litre of water sampled (n=3) for each MP type measured in the study. *** - indicate statistical significance of Two sample T-test (alpha=0.05).

### Host-Associated Microbiome Composition in Daphnia Linked to Environmental Variables

PERMANOVA showed a significant difference between the composition of bacterioplankton and *Daphnia* microbiome communities assessed with 16S rRNA sequencing (p < 0.01, betadisper > 0.05) (**Figure 3A**). We compared richness (Chao1, Shannon) and evenness (Pielou), finding bacterioplankton more diverse and even (p < 0.01 for all indices) than the field *Daphnia* microbiomes (**Figure 3B**).

**Figure 3.**
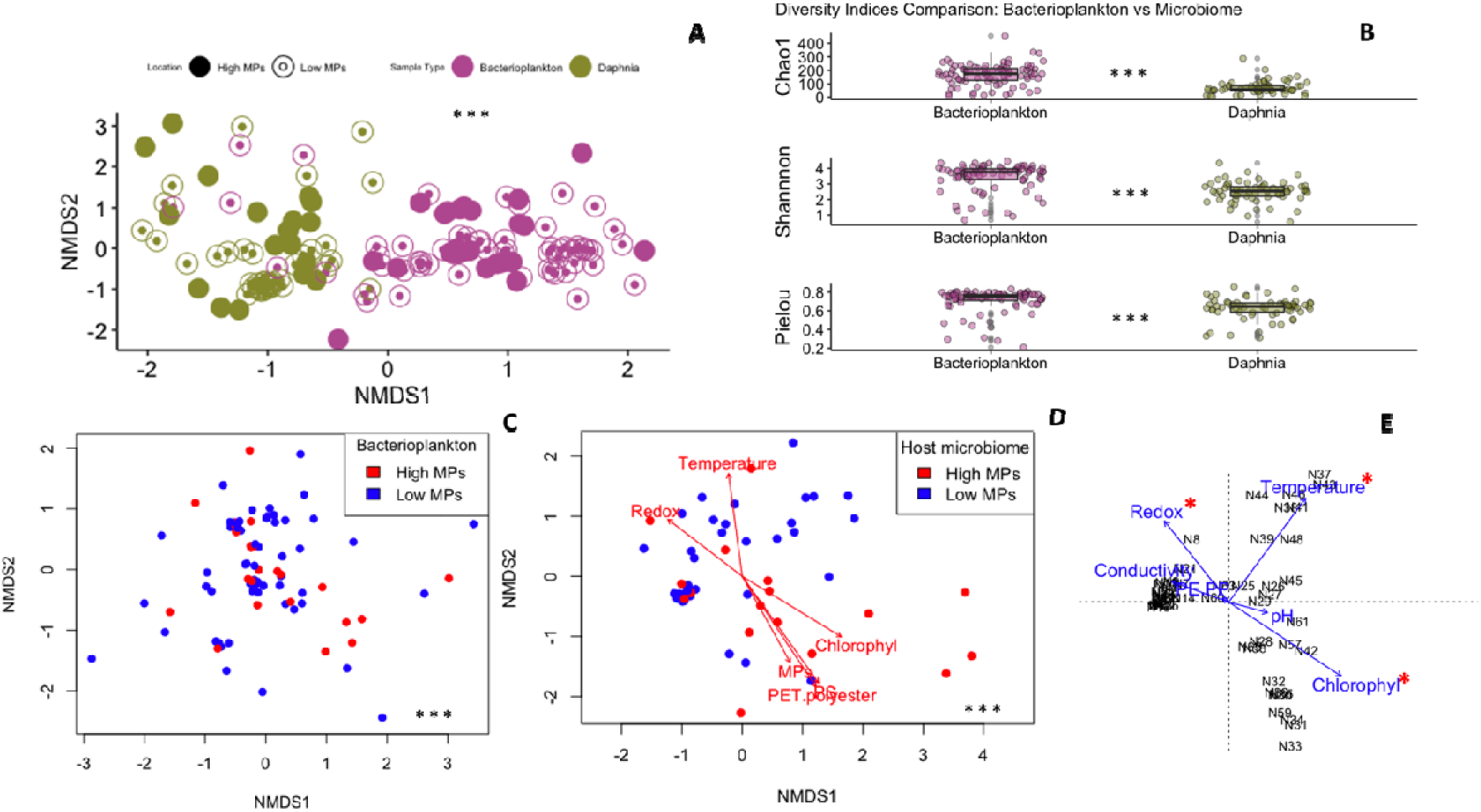
16S rRNA marker gene analysis of host-associated microbiome and bacterioplankton communities. **A.** NMDS ordination showing Bray-Curtis dissimilarities between bacterioplankton (purple) and host microbiome (green) samples and shapes depicting the type of the pond (high vs low MP). **B.** Comparison of diversity indices between bacterioplankton and *Daphnia* microbiome samples. **C.** NMDS ordination of bacterioplankton samples for high-MP (red) versus low-MP (blue) ponds with no significant environmental (envfit) factors. **D**. NMDS ordination of *Daphnia* host microbiome with significant envfit factors fitted to the sample coordinates and scaled according to the strength of correlation. **E**. Constrained ordination of db-RDA scores for *Daphnia* microbiome with significant predictors marked with a red star. *** - indicate statistical significance of PERMANOVA (**A, C, D**) and Welch Two-sample t-test (**B**) at alpha of 0.05.

Assessing each sample type separately, PERMANOVA indicated significant differences in community composition between high and low MP ponds (p < 0.01, betadisper > 0.05) for both bacterioplankton and host microbiome (**Figures 3C**, **3D**).

To assess the impact of environmental variables on community composition, the former were regressed onto PCoA axes with envfit, which showed a significant correlation of the host community composition with PET/polyester, polystyrene, total MP, chlorophyll, redox, and temperature (highest R² = 0.319) (Figure 3D). When accounting for interactions using db-RDA, only redox, temperature, and chlorophyll were significant predictors with a combined adjusted R^2^ of 0.21 (**Figure 3E)**.

No significant differences in richness and evenness were found between high and low MP samples for either community (**Supplementary Figure 2**).

### Increased abundance of PET catabolic genes and ARGs in microbiomes sampled from MP-rich ponds

We assessed whether the functional profiles of host microbiome and bacterioplankton samples are significantly different. The PCoA demonstrated a distinct partitioning between microbiome and bacterioplankton samples along the principal components (**Figure 4**) and a significant difference between low and high MP catabolic gene composition for microbiome samples (PERMANOVA, p-value < 0.05, betadisper > 0.05). ARG have been correlated with plastic pollution in previous studies (Vlaanderen *et al*. 2023), so we quantified their abundance in the host-associated microbiome samples. The analysis of average resistome abundances revealed a statistically significant elevation in ARGs within the microbiome of high MP ponds compared to those from low MP ponds, as depicted in **Figure 5** (Wilcoxon rank sum, p < 0.05). Furthermore, we quantified the genes encoding for degradation pathway of the PET polymer, including the first two steps: PET hydrolase (PETase) [EC:3.1.1.101] and mono(ethylene terephthalate) hydrolase (MHETase) [EC:3.1.1.102]. The results are presented as a heatmap in **Figure 6** showing mean abundances for each KO enzyme. Gene homologues encoding enzymes known to be involved in bacterial PET metabolism were only found in the high MP pond group, including the genes encoding the initial depolymerase PETase (as well as the MHETase).

**Figure 4.**
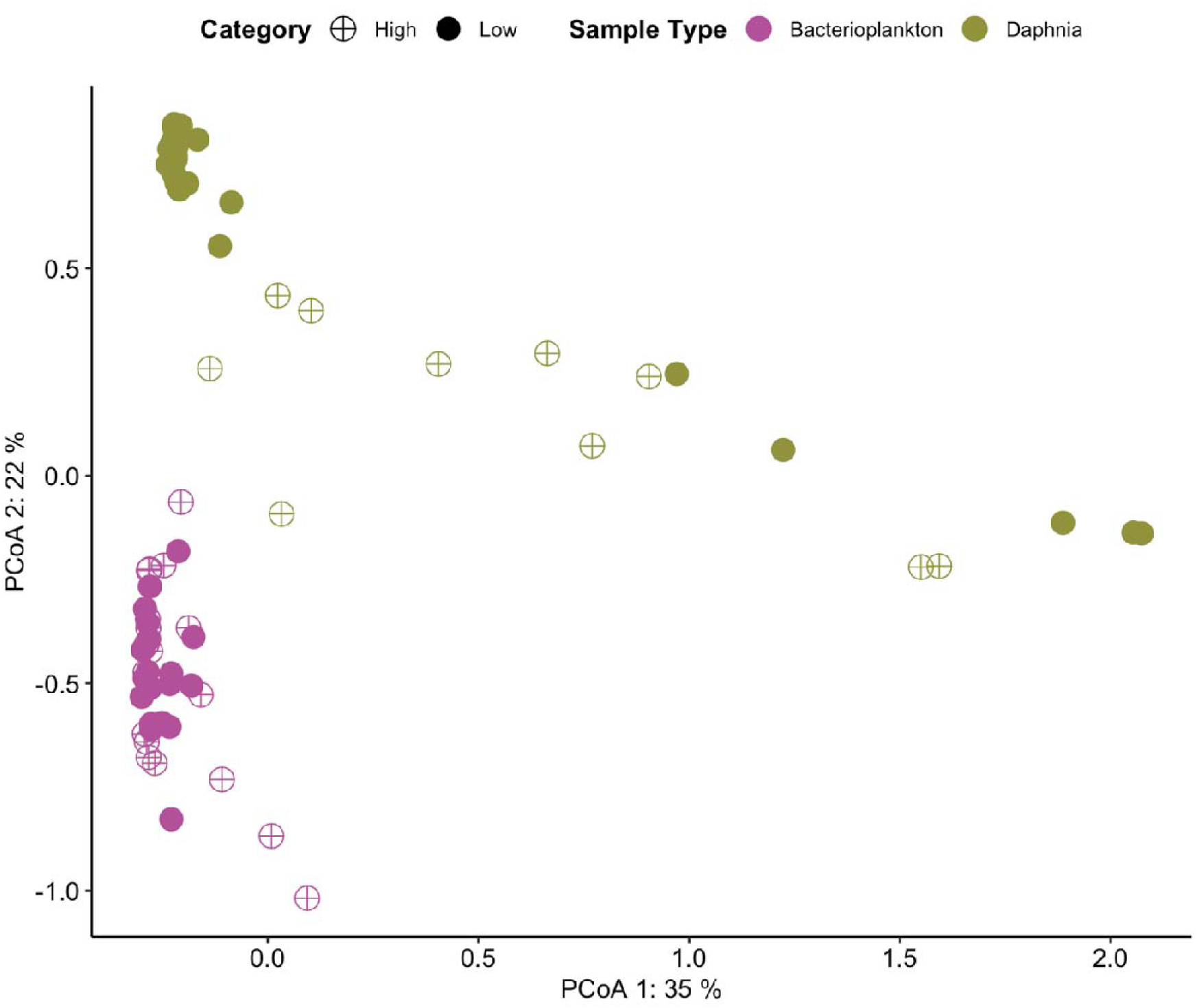
Difference in gene abundance between bacterioplankton and host microbiome. PCoA with Bray-Curtis dissimilarities between metagenome samples based on the rarefied abundance of gene functions (COGs). *Daphnia* metagenome samples are marked in green and bacterioplankton metagenomes in purple. Pond type according to high MP vs. low MP concentration, is indicated by ‘filled’ versus ‘cross’ circle symbols.

**Figure 5.**
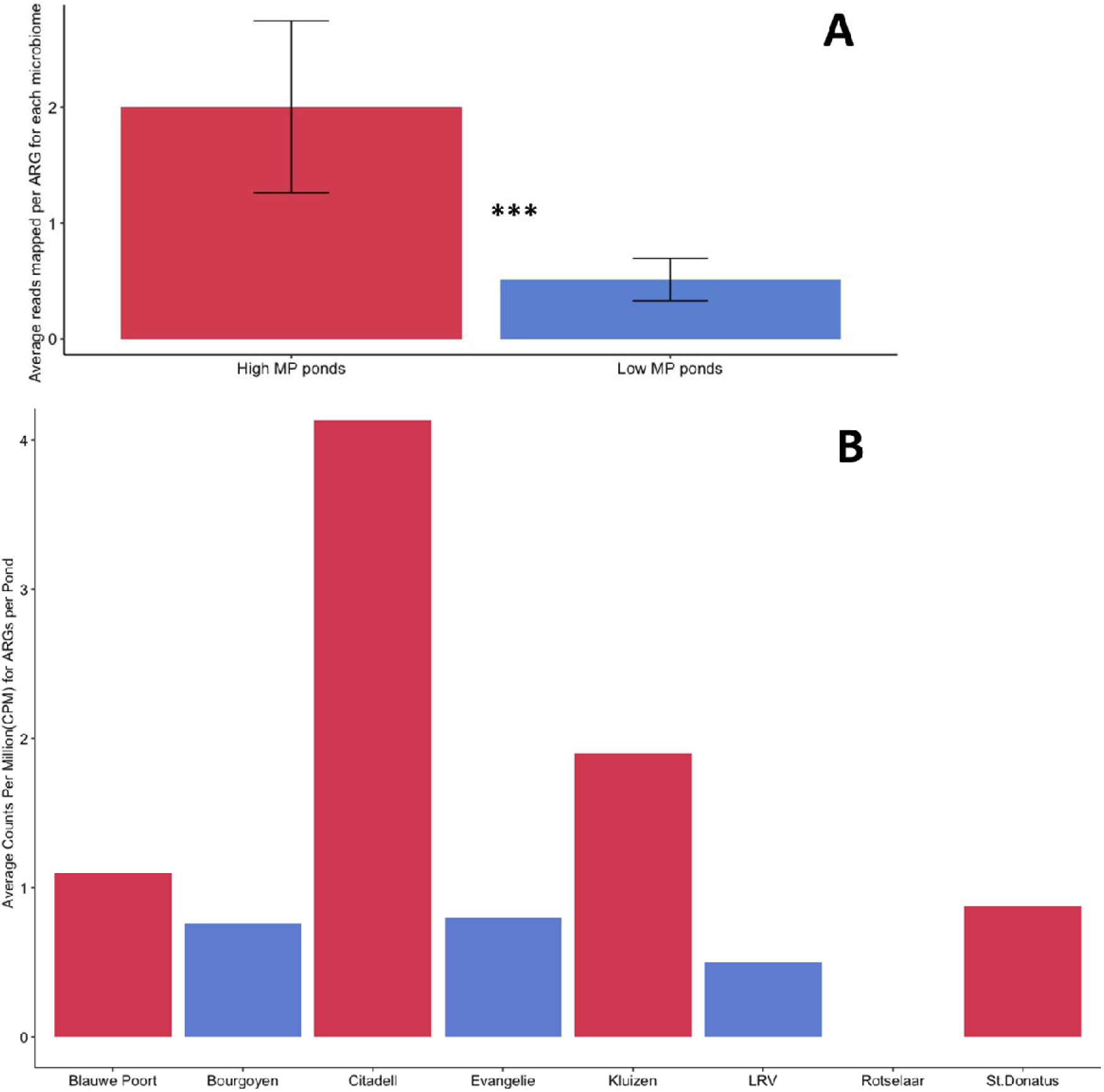
Difference in gene abundance between bacterioplankton and host microbiome and average abundance of ARG in host microbiome over the different ponds sampled (N=8). **A.** Bar chart showing abundance of ARG as average counts per million (CPM) per pond category. *** - indicates significant Wilcoxon rank sum at alpha of 0.05. **B.** Bar chart showing abundance of ARG as average counts per million (CPM) per individual pond. Colour red categorises the pond as “High MP” and blue as “Low MP”.

**Figure 6.**
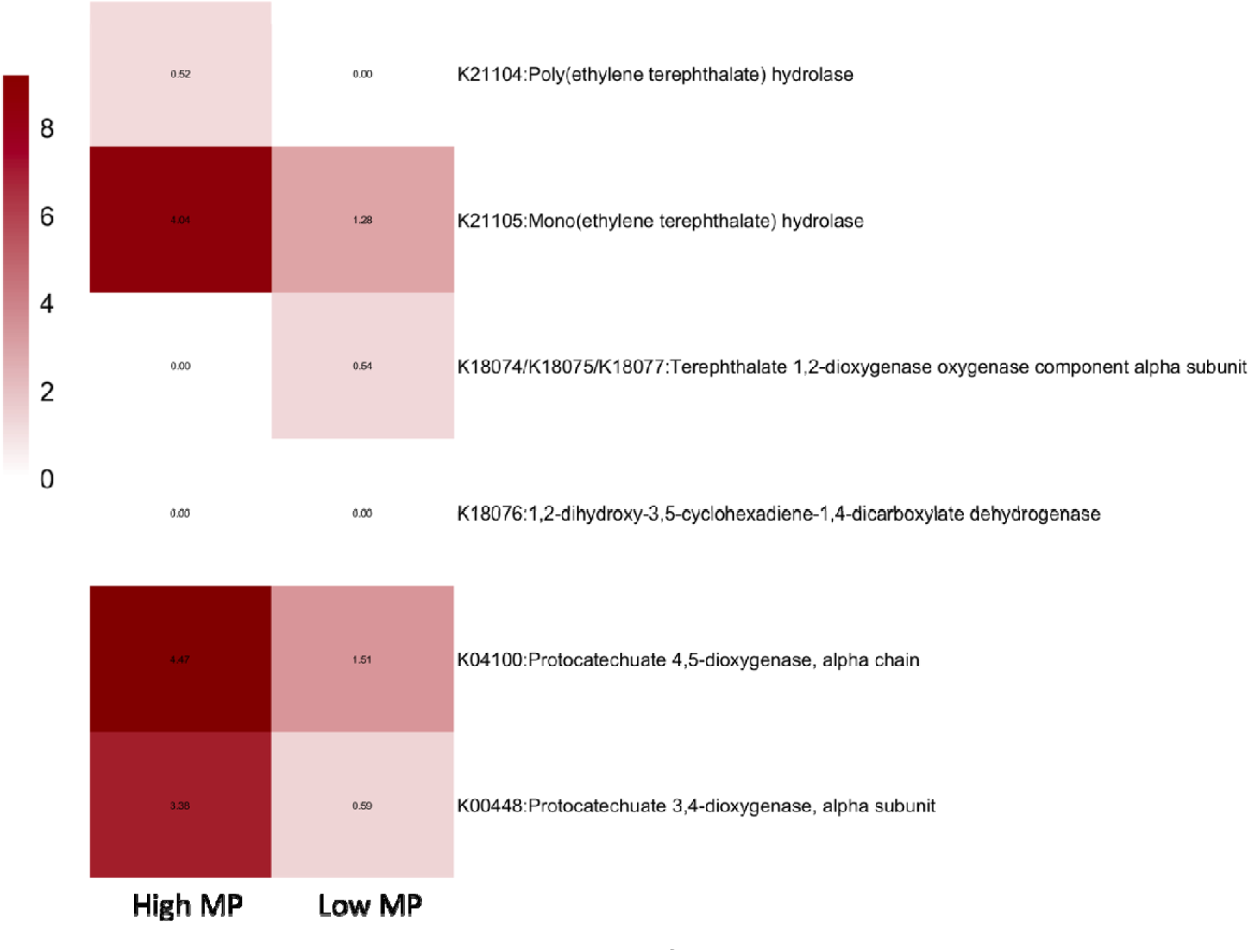
Heatmap showing abundances of enzymes involved in the PET degradation in host microbiome samples from high versus low MP ponds. The abundances are expressed as square roots of average CPM (Counts per million) reads mapped to KOs involved in the biodegradation of PET. The proteins were mapped to KOs using KEGG Mapper version 5.1.

### Bacterioplankton inoculum is the most important factor influencing microbiome composition of Daphnia in presence of MP

The *in situ* analysis showed differences in microbiome between high and low MP freshwaters with the high MP communities showing adaptation to MP pollution for instance by displaying PET catabolic genes. To examine the effects of MP exposure on the *Daphnia* composition in a controlled environment with previously adapted bacterioplankton inoculum, we conducted a 23-day exposure experiment. Specifically, different *Daphnia magna* genotypes were exposed to high MP concentrations with bacterioplankton inocula from either De-Gavers (DG) and Blauwe Poort (BP) as a low and high MP pond, respectively (**Figure 1E**). Treatment groups included three MP fibres (Nylon, PET, PLA), three *Daphnia* genotypes (KNO15.04, F, BH), and two water sources (BP and DG). We compared *Daphnia* microbiome composition before and after the experiment. PERMANOVA revealed significant changes (p < 0.05), likely due to bacterioplankton uptake. Post-experiment microbiomes differed significantly from surrounding bacterioplankton (p < 0.05) and between jars with and without *Daphnia* (p < 0.05), highlighting exchange between *Daphnia* microbiome and bacterioplankton in the presence of MP. No significant difference was found between MP-exposed and control microbiomes (p > 0.05). A multi-factor PERMANOVA indicated only pond bacterioplankton inoculum significantly affected post-exposure microbiomes (p < 0.05). For BP-inoculated samples, *Daphnia* development stage (juvenile vs adult) significantly affected microbiomes (p < 0.01), but no significant effects were found for DG-inoculated samples.

### Taxa associated to microplastics

We conducted a network analysis on DG and BP-inoculated *Daphnia* microbiome communities from the exposure experiment to uncover significant taxon-taxon and taxon-MP associations (**Supplementary table 1**). For the BP community, the genus *Prosthecobacter* was positively associated with PET and the order *Planctomycetales* and the genus *Gemmobacter*, positively with Nylon. We traced the origin of each taxon to either pre-treatment (taxa from *Daphnia* core microbiome), post-treatment (taxa derived from the bacterioplankton inoculum), or both. *Prosthecobacter* was identified both in the pre-treatment microbiome and BP inocula whereas *Planctomycetales* and *Gemmobacter* were unique to the BP inoculum. The low-MP inoculum (De Gavers) was not associated with any plastic variable.

### MP and bacterioplankton inoculum influence *Daphnia magna* survival and growth

*Daphnia* survival and body size were tracked throughout the experiment. Kaplan-Meier survival curves (**Figure 8A**) showed differing survival probabilities in response to MP fibres (Nylon, PET, PLA, control). A significant decline in survival was observed after 20 days, particularly in the PLA-treated groups. We examined the combined effects of fiber type and bacterioplankton inoculum (Fiber x Pond) on *Daphnia* survival (**Figure 8B**). At the end of the experiment, a sharp decline was observed in all groups, most notably in the DG x control group, where survival fell below 25%. Most *Daphnia* individuals survived in groups inoculated with high-MP bacterioplankton (BP x PET, BP x control, BP x PLA, BP x Nylon) while survival was lowest in groups with low-MP bacterioplankton (DG x Control, DG x PLA, DG x PET, DG x Nylon). Cox proportional hazards analysis indicated that *Daphnia* inoculated with DG pond bacterioplankton had a 24.78% higher risk of death compared to those inoculated with BP bacterioplankton. PLA fibers were associated with a 47.51% higher risk of death than other fibers and control groups. Analysis of the day of the first brood, an important reproductive parameter, showed that *Daphnia* from the DG pond reached this stage significantly earlier than those from BP ponds, suggesting an accelerated life cycle in response to local environmental stress for DG-inoculated individuals.

**Figure 7.**
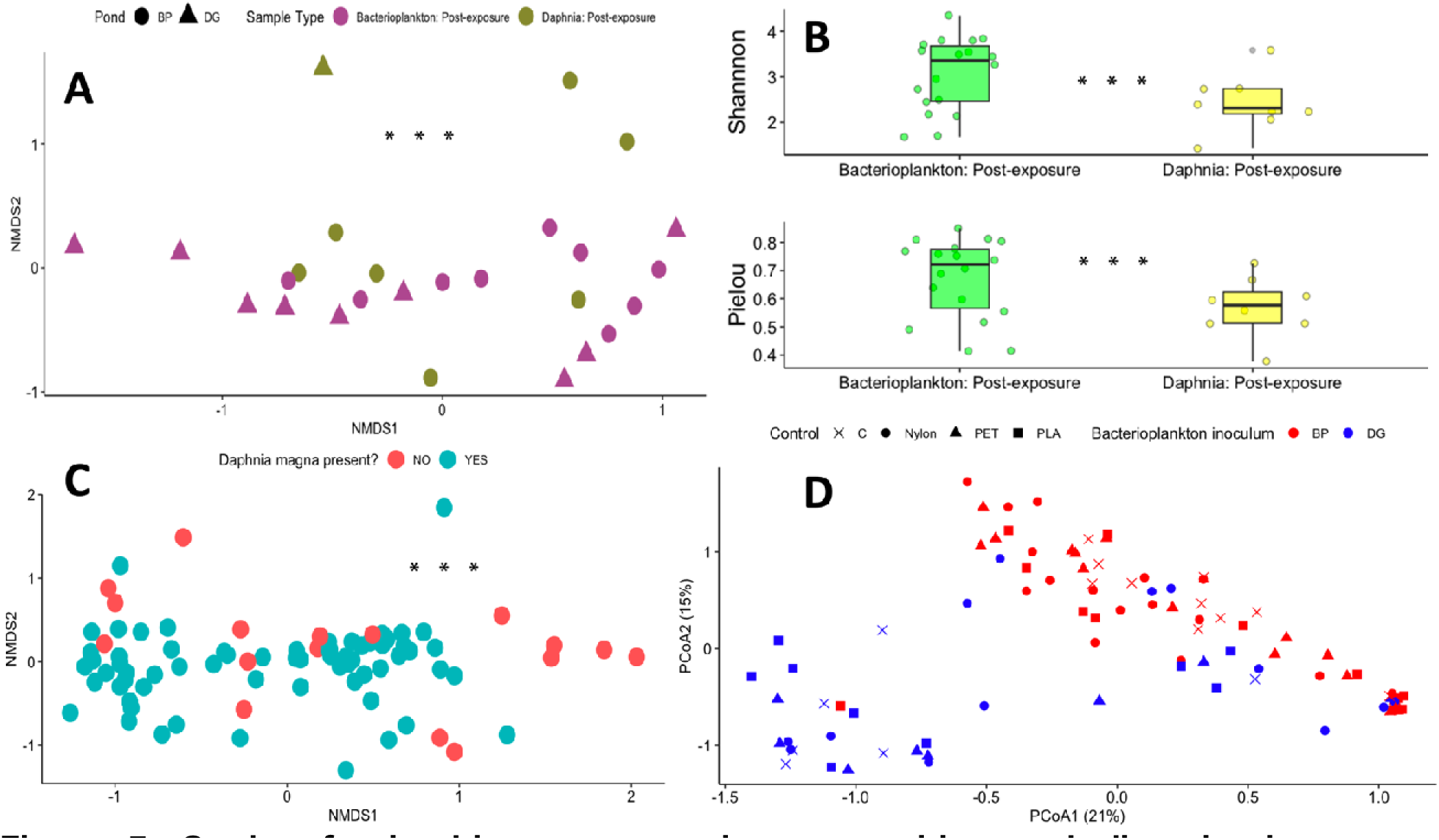
Study of microbiome community composition and diversity in bacterioplankton and *Daphnia* microbiome samples. **A.** PCoA ordination of Bray-Curtis dissimilarities for samples after the exposure experiment stratified by type (Bacterioplankton, *Daphnia magna* microbiome) and pond inoculum source (Blauwe-Poort, De Gavers). **B.** Richness and evenness indices for Bacterioplankton and *Daphnia magna* microbiome communities after the exposure. **C.** PCoA ordination of Bray-Curtis dissimilarities for bacterioplankton samples co-cultured with *Daphnia magna* (blue) and without (red). **D.** PCoA ordination of Bray-Curtis dissimilarities for *Daphnia magna* microbiome samples after 23 days of experiment. BP pond inoculated samples are shown in red and DG-inoculated in blue. The MP treatment is represented as a shape. *** - indicates significant PERMANOVA for ordination plots and significant Wilcoxon rank sum for richness indices. Alpha was set at 0.05 for all tests.

**Figure 8.**
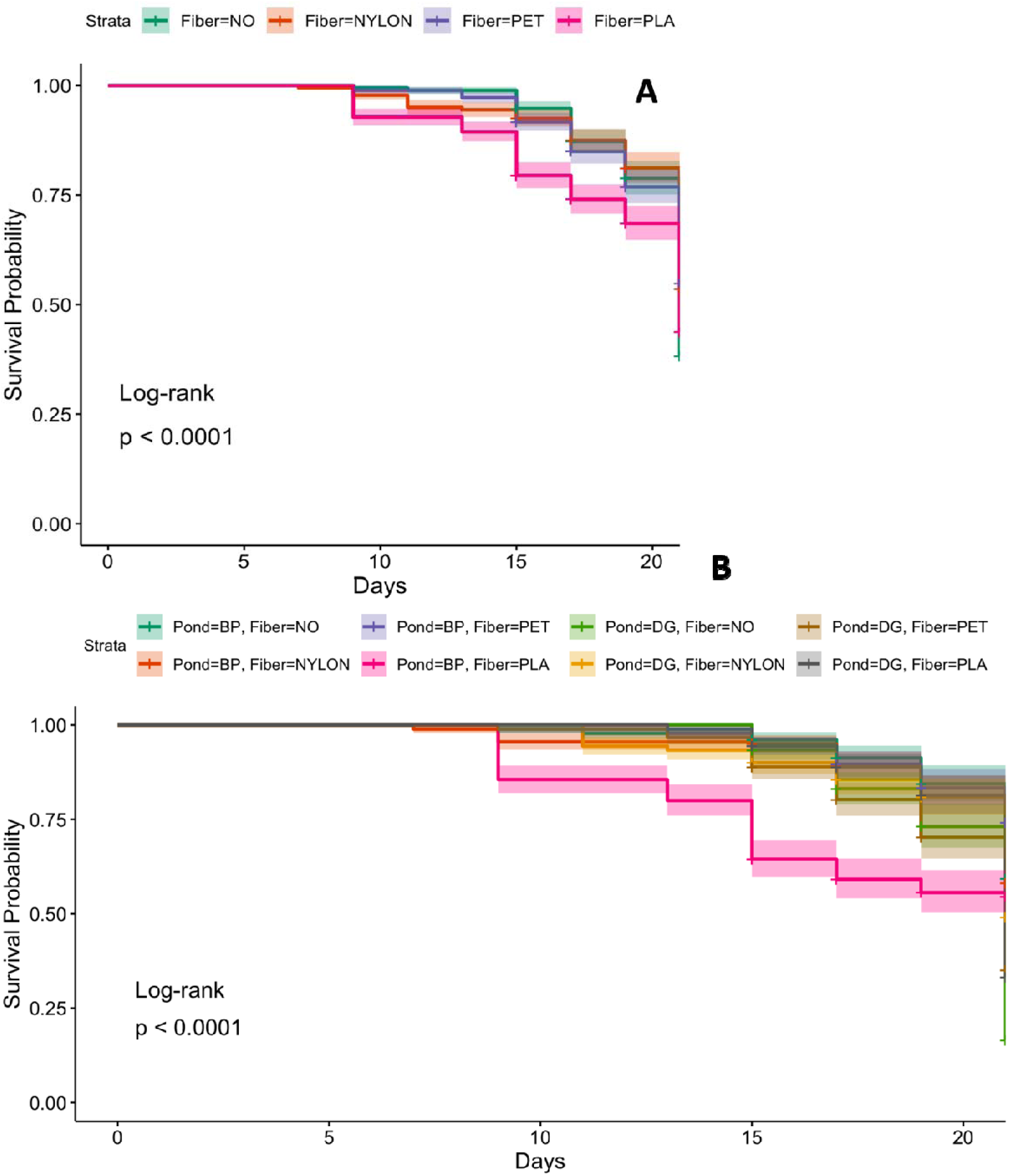
Kaplan-Meier survival curves showing survival of *Daphnia magna* population over 23 days exposure experiment. **A.** Survival curves stratified by type of fibre used in the study (Fiber=NO represents a control) . **B.** Survival curves with combinations of Fiber x Pond. The significance was tested with log-rank tests.

## DISCUSSION

### Microplastic Profiles in Urban Water Bodies: Sources, Composition, and Accumulation

This study highlights the impact of urbanisation on MP distribution in aquatic environments, as shown by significantly higher MP concentrations in city pond sediments and water compared to natural lakes. Our findings align with previous research indicating that urban water bodies serve as substantial sinks for MP due to elevated anthropogenic activities (Nava et al., 2023; Xiong et al.,2018; Scherer et al., 2020; Liu et al., 2019). Furthermore, we observed a discrepancy in the concentration and composition of MP between sediment and water samples in the urban setting. Specifically, sediment MP were predominantly composed of PP and PE, whereas water samples were enriched with polyesters, including PET. Plastic fragments will deposit at the bottom sediment if their density surpasses the water (>1 g/cm^3^) e.g. nylon, polyvinyl chloride, acrylic (Corcoran, 2015). On the other hand, low-density polymers PE/PP (0.89-0.97 g/cm³) have been commonly found to be abundant in freshwater sediments (Fan et al., 2019; Zhang et al., 2015; Sruthy & Ramasamy, 2017;Rodrigues et al., 2018; Zhou et al., 2018). Although these polymers are likely to float, there are several reasons for their increased sedimentation such as biofouling (Morét-Ferguson et al., 2010, Lobelle & Cunliffe, 2011) or injection with mineral fillers during the manufacturing to improve their structural properties (Corcoran, 2015). These processes will also affect the accumulation rate of MP across the layers. Notably, we observed a higher abundance of MP in sediment than in the water column. Benthic MP will biodegrade slower due to a decreased photooxidative effect and thus, its persistence will be higher than free-floating MP particles (Andrady et al., 2011). While PE/PP derive from fragmentation of discarded plastic waste, polyesters enter aquatic environments via a different route. A large proportion of polyesters are released into freshwater ecosystems from wastewater effluent due to industrial and household textile washing (Galvao et al., 2020; De Falco et al., 2020; Hernandez et al. 2017; Cai et al., 2020; Kelly et al., 2019). Despite their higher relative abundance in the water column, the actual concentration of polyester fibres is likely much higher in the tested ponds. This is due to their small size, typically in the range of 10-20 µm, which may have been missed given the 64 µm size threshold chosen for the study (Yang et al., 2023).

### Daphnia Microbiome Composition Reflects a Balance Between Host-Specific and Environmental Bacteria

Previous studies showed that *Daphnia* microbiome composition strongly depends on the diversity of bacteria in the surrounding bacterioplankton (Callens et al. 2020). Using 16S rRNA gene amplicon sequencing, we found a significant difference between the composition of the bacterioplankton community and the host microbiome in sampled ponds, irrespective of their contamination status (low MP vs high MP). Moreover, bacterioplankton was on average richer and more diverse than the related host microbiome. This finding is consistent with previous observations in *in vitro* experiments by Macke et al. (2020) and was also replicated in the MP exposure experiment showing a significantly altered *Daphnia magna* microbiome composition after 23 days of incubation while remaining distinct from the surrounding bacterioplankton. Interestingly, the composition of the bacterioplankton also varied depending on whether it was cultured with or without *Daphnia*. This suggests that *Daphnia* not only incorporates external bacteria but also maintains a core set of microbial species, leading to the emergence of a mixed community that comprises both the newly assimilated bacterioplankton and the native microbiome members (Callens et al. 2020; Freese and Schink 2011; Macke et al. 2017, 2020).

### Effects of MP on Daphnia microbiome community

Uptake of MP by *Daphnia* was shown in previous *in vitro* exposure studies (Elizalde-Velázquez *et al*., 2020; Trotter *et al*. 2021; Jemec *et al*. 2016; Liu *et al*., 2020). In this study, we compared host microbiome and bacterioplankton communities sampled from ponds burdened with MP pollution with those of lower MP exposure. Significant differences in microbial community composition were found between the two environments although their diversity did not differ significantly. MP has been shown before to alter microbial community composition in sediments (Seeley et al., 2020; Li et al., 2020; Sun et al., 2022) However, their effects on host-associated microbiomes, particularly within freshwater environments, remain largely unexplored. Considering the complexity of aquatic and host-associated ecosystems, the presence of MP is unlikely to affect microbial community structure without consideration for other interacting factors including environmental variables. Indeed, urban ponds, categorised in this study as high-MP environments, can differ significantly from non-urban lakes in several aspects including among others nutrient levels, physical disturbances, chemical contamination, and habitat structure (Paul et al., 2001). Our findings indicate that *Daphnia* microbiome communities in urban ponds with high MP pollution were correlated with chlorophyll concentration, which is a proxy for the amount of algae, including cyanobacteria. Urban environments often contribute to higher nutrient loads (e.g., from runoff containing fertilisers and pollutants from residential and industrial areas), which can exacerbate algal blooms including *Cyanobacteria* which thrive in nutrient-rich conditions (Grogan et al., 2023). Exposure to cyanobacteria and their toxins has been shown to disrupt the balance of the gut microbiome in *Daphnia* (Macke et al. 2017). *Daphnia* microbiome communities in low-MP ponds were more strongly associated with temperature and oxygen availability indicating that in the absence of additional environmental stress, temperature and oxygen may play a more significant role in shaping microbial communities. Oxygen depletion has also been shown to structure *Daphnia* microbiome communities in experiments (Coone et al. 2023). Given our observations in the field study, we aimed to investigate the changes in the *Daphnia magna* microbiome community after exposure to MP while controlling for external variables. A 23-day *in-vitro* exposure of *Daphnia magna* to MP revealed no significant effects of MP fibres (PLA, PET, Nylon) on the community structure. However, a few taxa were associated with MP.

### Potential Plastisphere: Taxonomic and functional enrichment in presence of MP

Our network analysis of exposure study samples revealed that a taxon classified as *Prosthecobacter*, a member of both *Daphnia* core microbiome and bacterioplankton inoculum community, was linked to PET. This genus is known to thrive in low-nutrient environments and was observed to be enriched in the presence of other plastic polymers including the deep sea plastisphere (Kelly et al., 2022), PET film in stream water (Rummel et al., 2021) and cellulose acetate polymer surface in a brackish marine environment (Eroen-Rasimus et al., 2022). However, this is the first time that its association with PET is reported for freshwater. In addition, two taxa belonging to the Gemmobacter genus and *Plantomycetales* order were linked to Nylon. *Gemmobacter* belongs to the *Rhodobacteraceae* family, which is commonly found on plastic (Zettler et al., 2013; Bryant et al., 2016) and has been recognized as an initial coloniser during biofilm formation (Elifantz et al., 2013). Moreover, it was observed in biofilms developing on polypropylene films in a Hungarian lake (Szabó et al., 2020) Not much is known about the *Plantomycetales* order in relation to plastic colonisation or degradation except that they were identified together with *Gemmobacter* as members of the freshwater biofilms forming on polypropylene films in the study of Szabó et al. (2020). Moreover, they were found implicated in biofilm formation on various macroalgae (Bengtsson & Øvreås, 2010; Burke et al., 2011; Lachnit et al., 2011). No taxa associated with the PLA were identified.

By applying metagenome shotgun sequencing to field samples, we quantified specific functional genes in *Daphnia* microbiomes that could be associated with MP presence, including ARGs and hydrolytic enzymes involved in polyester biodegradation. Recently, plastispheres have been identified as hotspots for ARGs (Zhu et al., 2022; Vlaanderen et al., 2023; Zadjelovic et al., 2023) and hence their higher abundances in the host microbiomes from high-MP ponds was not unexpected. Since high-MP ponds are close to human activity hubs, they are at greater risk of antibiotic contamination through sewage runoff. Furthermore, the increase of ARGs within host microbiomes in high-MP ponds supports the hypothesis that MP may provide a route for human-derived antibiotics and ARGs to enter and establish within aquatic host microbiomes (Zhao et al., 2024; Chaturvedi et al., 2023). Interestingly, we found that the host microbiome community from a high-MP environment was enriched in genes encoding PET catabolism. Polyesters are common in freshwater (Li et al., 2019) and were also the most abundant MP type in the urban pond waters in the current study. Therefore, these findings support the hypothesis that biofilm-associated microbes may become part of MP-grazing microbiomes as evidenced by enrichment of plastic/biofilm associated taxa in the microbiome community as well as increase in genes involved in polyester degradation and antibiotic resistance. However, additional research is required to confirm these interactions and fully understand the mechanisms by which MP-associated biofilms influence host microbiomes in freshwater ecosystems.

### Effects of MP on Daphnia magna survival

Finally, we assessed whether bacterioplankton pre-exposed to different levels of MP, *Daphnia* genotype and MP type affected the survival of *Daphnia magna.* The host genotype did not matter, but *Daphnia* exposed to PLA fibres exhibited accelerated mortality and a 48% increased risk of death compared to *Daphnia* exposed to other MP types. PLA particles used in the study measured 50×50 µm, as opposed to 50×10µm for PET and Nylon. *Daphnia* ingests larger MP particles at a greater rate than smaller particles (Wang & Wang, 2023), suggesting that the PLA particles used in the study might have accumulated faster than the two other MP in the *Daphnia* digestive tract, potentially obstructing it and hindering proper digestion and nutrient absorption, ultimately leading to starvation. Another significant factor is the pond inoculum effect: *Daphnia* cultured with bacterioplankton derived from the MP-rich BP pond had the highest survival rates and those cultured with the MP-poor DG pond were 25% more at risk of death. Thus, bacterioplankton pre-exposed to high levels of MP appears more beneficial to *Daphnia* survival. We assume that the high-MP environment selects for taxa linked to MP degradation, which are then transferred to and become part of *Daphnia’s* microbiome. However, further studies are required to confirm this hypothesis and to unravel the mechanisms behind the observed impact on *Daphnia* survival.

### Conclusions

In conclusion, our study highlights consequences associated with urbanisation for aquatic ecosystems such as higher MP and ARG burden. Moreover, our combined *in situ* and *in vitro* approach demonstrates the enrichment of plastisphere-associated taxa and/or plastic-degradation functions in the *Daphnia* microbiome as a result of MP presence. Our research opens avenues to further explore the adaptive mechanisms of aquatic microbiomes and acclimatisation of keystone invertebrates in response to MP pollution, emphasising the complex interactions within ecosystems and the potential long-term impacts of urbanisation.

### Data availability statement

#### Reads and metadata are available at PRJNA1122644

Field’s whole genome shotgun reads accessions: SAMN41788498-526 Field’s amplicon 16 rRNA Miseq reads accessions: SAMN41798624-771 MP exposure amplicon 16 rRNA Miseq reads accessions: SAMN41876284-514

#### Scripts and metadata

github.com/AMK06-1993/Effects-of-microplastics-on-Daphnia-associated-microbiomes-in-situ-and-in-vitro

### Authors’ contributions

K.F. and E.D conceived and designed the experiments. A.K., N.P., I.V., J.L. and B.V.D.M performed the experiments. A.K. analyzed the data. W.T contributed reagents. D.S. provided NPOCs measurments. D.D. performed MP quantification, A.K., K.F. and E.D. wrote the manuscript. A.K., K.F., E.D., K.M and D.S reviewed and edited the manuscript. All authors read and approved the final manuscript.

## Acknowledgements

We would like to thank Annelien Verfaillie and Wouter Bossuyt from Genomics Core (Leuven) for helping us optimise the sequencing protocol.

## References

1. Aljaibachi R, Laird WB, Stevens F, Callaghan A. Impacts of polystyrene microplastics on *Daphnia magna*: A laboratory and a mesocosm study. Science of The Total Environment. 2020;705:135800. 10.1016/j.scitotenv.2019.135800

2. Altschul SF, Gish W, Miller W, Myers EW, Lipman DJ. Basic local alignment search tool. Journal of Molecular Biology. 1990;215(3):403–410. 10.1016/S0022-2836(05)80360-2

3. Andrady AL. Microplastics in the marine environment. Mar Pollut Bull. 2011;62(8):1596–605. doi: 10.1016/j.marpolbul.2011.05.030. PMID: 21742351.

4. Andrews S. FastQC: A Quality Control Tool for High Throughput Sequence Data. Available online at: http://www.bioinformatics.babraham.ac.uk/projects/fastqc/

5. Barata C, Campos B, Rivetti C, LeBlanc GA, Eytcheson S, McKnight S, et al. Validation of a two-generational reproduction test in *Daphnia magna*: An interlaboratory exercise. Science of the Total Environment. 2017;579:1073–1083. 10.1016/j.scitotenv.2016.11.066

6. Bengtsson MM, Øvreås L. Planctomycetes dominate biofilms on surfaces of the kelp *Laminaria hyperborea*. BMC Microbiol. 2010;10:261. 10.1186/1471-2180-10-261

7. Borrelle SB, Ringma J, Law KL, Monnahan CC, Lebreton L, McGivern A, et al. Predicted Growth in Plastic Waste Exceeds Efforts to Mitigate Plastic Pollution. Science. 2020;369:1515–1518. doi: 10.1126/science.aba3656

8. Benjamini Y, Hochberg Y. Controlling the False Discovery Rate: A Practical and Powerful Approach to Multiple Testing. J R Stat Soc Series B Methodol. 1995;57(1):289–300. Available from: http://www.jstor.org/stable/2346101

9. Bryant JA, Clemente TM, Viviani DA, Fong AA, Thomas KA, Kemp P, et al. Diversity and activity of communities inhabiting plastic debris in the North Pacific Gyre. MSystems. 2016;1(3). 10.1128/mSystems.00024-16

10. Cai Y, Yang T, Mitrano DM, Heuberger M, Hufenus R, Nowack B. Systematic Study of Microplastic Fiber Release from 12 Different Polyester Textiles during Washing. Environ Sci Technol. 2020;54:4847–4855. [Google Scholar] [CrossRef]

11. Callahan BJ, McMurdie PJ, Rosen MJ, Han AW, Johnson AJA, Holmes SP. DADA2: High-resolution sample inference from Illumina amplicon data. Nature Methods. 2016;13(7):581–583. 10.1038/nmeth.3869

12. Callens M, De Meester L, Muylaert K, Mukherjee S, Decaestecker E. The bacterioplankton community composition and a host genotype dependent occurrence of taxa shape the *Daphnia magna* gut bacterial community. FEMS Microbiol Ecol. 2020;96(8).

13. Callens M, Macke E, Muylaert K, et al. Food availability affects the strength of mutualistic host–microbiota interactions in Daphnia magna. ISME J. 2016;10:911–920. Available from: 10.1038/ismej.2015.166

14. Chaturvedi A, Li X, Dhandapani V, Marshall H, Kissane S, Cuenca-Cambronero M, Asole G, Calvet F, Ruiz-Romero M, Marangio P, Guigó R, Rago D, Mirbahai L, Eastwood N, Colbourne JK, Zhou J, Mallon E, Orsini L. The hologenome of Daphnia magna reveals possible DNA methylation and microbiome-mediated evolution of the host genome. Nucleic Acids Res. 2023 Oct 13;51(18):9785–9803. doi: 10.1093/nar/gkad685.

15. Cole M. A novel method for preparing microplastic fibers. Scientific Reports. 2016;6:34519. 10.1038/srep34519

16. Coone M, Vanoverberghe I, Houwenhuyse S, Verslype C, Decaestecker E. The effect of hypoxia on *Daphnia magna* performance and its associated microbial and bacterioplankton community: A scope for phenotypic plasticity and microbiome community interactions upon environmental stress? Frontiers in Ecology and Evolution. 2023;11. 10.3389/fevo.2023.1131203

17. Cooper RO, Cressler CE. Characterization of key bacterial species in the *Daphnia magna* microbiota using shotgun metagenomics. Sci Rep. 2020;10:652. 10.1038/s41598-019-57367-x

18. Corcoran PL. Benthic plastic debris in marine and fresh water environments. Environmental Science: Processes & Impacts. 2015;17(8):1363–1369. doi:10.1039/c5em00188a.

19. Cui R, Kim SW, An YJ. Polystyrene nanoplastics inhibit reproduction and induce abnormal embryonic development in the freshwater crustacean *Daphnia galeata*. Sci Rep. 2017;7:12095. 10.1038/s41598-017-12299-2

20. De Falco F, Cocca MC, Avella M, Thompson RC. Microfiber Release to Water, Via Laundering, and to Air, via Everyday Use: A Comparison between Polyester Clothing with Differing Textile Parameters. Environ Sci Technol. 2020;54:3288–3296. [Google Scholar] [CrossRef] [PubMed]

21. De Felice B, Sugni M, Casati L, Parolini M. Molecular, biochemical and behavioral responses of *Daphnia magna* under long-term exposure to polystyrene nanoplastics. Environment International. 2022;164:107264. 10.1016/j.envint.2022.107264

22. Dillies MA, Rau A, Aubert J, Hennequet-Antier C, Jeanmougin M, Servant N, et al. A comprehensive evaluation of normalization methods for Illumina high-throughput RNA sequencing data analysis. Briefings in Bioinformatics. 2013;14(6):671–683. 10.1093/bib/bbs046

23. Eckert EM, Pernthaler J. Bacterial epibionts of *Daphnia*: a potential route for the transfer of dissolved organic carbon in freshwater food webs. ISME J. 2014;8:1808–1819.

24. Elifantz H, Horn G, Ayon M, Cohen Y, Minz D. Rhodobacteraceae are the key members of the microbial community of the initial biofilm formed in Eastern Mediterranean Coastal Seawater. FEMS Microbiol Ecol. 2013;85(2):348–357. 10.1111/1574-6941.12122

25. Elizalde-Velázquez A, Carcano AM, Crago J, Green MJ, Shah SA, Cañas-Carrell JE. Translocation, trophic transfer, accumulation and depuration of polystyrene microplastics in *Daphnia magna* and *Pimephales promelas*. Environmental Pollution. 2020;259:113937. 10.1016/j.envpol.2020.113937

26. Eronen-Rasimus EL, Näkki PP, Kaartokallio HP. Degradation rates and bacterial community compositions vary among commonly used bioplastic materials in a brackish marine environment. Environmental Science & Technology. 2022;56(22):15760–15769. 10.1021/acs.est.2c06280

27. Fan Y, Zheng K, Zhu Z, Guangshi C, Peng X. Distribution, sedimentary record, and persistence of microplastics in the Pearl River catchment, China. Environ Pollut. 2019;251:862–870.

28. Frankel-Bricker J, Song MJ, Benner MJ, et al. Variation in the Microbiota Associated with *Daphnia magna* Across Genotypes, Populations, and Temperature. Microb Ecol. 2020;79:731–742. 10.1007/s00248-019-01412-9

29. Freese HM, Schink B. Composition and stability of the microbial community inside the digestive tract of the aquatic crustacean *Daphnia magna*. Microb Ecol. 2011;62(4):882–894. 10.1007/s00248-011-9886-8

30. Galvão A, Aleixo M, De Pablo H, Lopes C, Raimundo J. Microplastics in wastewater: Microfiber emissions from common household laundry. Environ. Sci. Pollut. Res. 2020;27:26643–26649. [Google Scholar] [CrossRef] [PubMed]

31. Grogan AE, Alves-de-Souza C, Cahoon LB, Mallin MA. Harmful Algal Blooms: A Prolific Issue in Urban Stormwater Ponds. Water. 2023;15(13):2436. Available from: 10.3390/w15132436

32. Gökçe D, Köytepe S, Özcan İ. Effects of nanoparticles on *Daphnia magna* population dynamics. Chemistry and Ecology. 2018;34(4):301–323. 10.1080/02757540.2018.1429418

33. Gorokhova E, Motiei A, El-Shehawy R. Understanding biofilm formation in ecotoxicological assays with natural and anthropogenic particulates. Frontiers in Microbiology. 2021;12:632947. 10.3389/fmicb.2021.632947

34. Howard SA, McCarthy RR. Modulating biofilm can potentiate activity of novel plastic-degrading enzymes. npj Biofilms and Microbiomes. 2023;9:72. 10.1038/s41522-023-00440-1

35. Huerta-Cepas J, Szklarczyk D, Heller D, Hernández-Plaza A, Forslund SK, Cook H, et al. eggNOG 5.0: a hierarchical, functionally and phylogenetically annotated orthology resource based on 5090 organisms and 2502 viruses. Nucleic Acids Research. 2019;47(D1). 10.1093/nar/gky1085

36. Houwenhuyse S, Stoks R, Mukherjee S, et al. Locally adapted gut microbiomes mediate host stress tolerance. ISME J. 2021;15:2401–2414. 10.1038/s41396-021-00940-y

37. Huang A, Cao S, Sun F, Wang L, Guo H, Ji R. Effects of nanoplastics and microplastics on toxicity, bioaccumulation, and environmental fate of phenanthrene in fresh water. Environmental Pollution. 2016;219:166–173. 10.1016/j.envpol.2016.10.061

38. Huerta-Cepas J, Szklarczyk D, Heller D, Hernández-Plaza A, Forslund SK, Cook H, et al. eggNOG 5.0: a hierarchical, functionally and phylogenetically annotated orthology resource based on 5090 organisms and 2502 viruses. Nucleic Acids Research. 2019;47(D1). 10.1093/nar/gky1085

39. Jaikumar G, Brun NR, Vijver MG, Bosker T. Reproductive toxicity of primary and secondary microplastics to three cladocerans during chronic exposure. Environmental Pollution. 2019;249:638–646. 10.1016/j.envpol.2019.03.085

40. Jemec A, Horvat P, Kunej U, Bele M, Kržan A. Uptake and effects of microplastic textile fibers on freshwater crustacean *Daphnia magna*. Environ Pollut. 2016 Dec;219:201–209. doi: 10.1016/j.envpol.2016.10.037. Epub 2016 Oct 28. PMID: 27814536.

41. Jia B, Raphenya AR, Alcock B, Waglechner N, Guo P, Tsang KK, et al. CARD 2017: expansion and model-centric curation of the comprehensive antibiotic resistance database. Nucleic Acids Research. 2017;45(D1). 10.1093/nar/gkw1004

42. Kanehisa M, Sato Y, Morishima K. BlastKOALA and GhostKOALA: KEGG tools for functional characterization of genome and metagenome sequences. Journal of Molecular Biology. 2015;428(4):726–731. 10.1016/j.jmb.2015.11.006

43. Kaur K, Reddy S, Barathe P, Oak U, Shriram V, Kharat SS, et al. Microplastic-associated pathogens and antimicrobial resistance in environment. Science of The Total Environment. 2022;813:151902. 10.1016/j.scitotenv.2021.151902

44. Kelly MR, Lant NJ, Kurr M, Burgess JG. Importance of Water-Volume on the Release of Microplastic Fibers from Laundry. Environ Sci Technol. 2019;53(20):11735–11744. doi: 10.1021/acs.est.9b03022.

45. Kelly MR, Whitworth P, Jamieson A, Burgess JG. Bacterial colonisation of plastic in the Rockall Trough, North-East Atlantic: An improved understanding of the deep-sea plastisphere. Environmental Pollution. 2022;305:119314. 10.1016/j.envpol.2022.119314

46. Lachnit T, Meske D, Wahl M, Harder T, Schmitz R. Epibacterial community patterns on marine macroalgae are host-specific but temporally variable. Environ Microbiol. 2011;13:655–665. doi: 10.1111/j.1462-2920.2010.02371.x

47. Langmead B, Salzberg SL. Fast gapped-read alignment with Bowtie 2. Nature Methods. 2012;9:357–359. 10.1038/nmeth.1923

48. Leo Lahti, Sudarshan Shetty et al. (Bioconductor, 2017). Tools for microbiome analysis in R. Microbiome package version 1.23.1. URL: http://microbiome.github.com/microbiome.

49. Li C, Busquets R, Campos LC. Assessment of microplastics in freshwater systems: A review. Sci Total Environ. 2020 Mar 10;707:135578. doi: 10.1016/j.scitotenv.2019.135578. Epub 2019 Nov 20. PMID: 31784176.

50. Li H, Handsaker B, Wysoker A, Fennell T, Ruan J, Homer N, et al. The Sequence Alignment/Map format and SAMtools. Bioinformatics. 2009;25(16):2078–2079. 10.1093/bioinformatics/btp352

51. Li Y, Xu Z, Liu H. Nutrient-imbalanced conditions shift the interplay between zooplankton and gut microbiota. BMC genomics. 2021;22:1–18.

52. Liao Y, Smyth GK, Shi W. featureCounts: an efficient general-purpose program for assigning sequence reads to genomic features. Bioinformatics. 2014;30(7):923–930. 10.1093/bioinformatics/btt656

53. Li Y, Yan N, Wong TY, Wang WX, Liu H. Interaction of antibacterial silver nanoparticles and microbiota-dependent holobionts revealed by metatranscriptomic analysis. Environ. Sci. Nano. 2019;6(11):3242–3255. 10.1039/c9en00587k

54. Li Y, Xu Z, Liu H. Nutrient-imbalanced conditions shift the interplay between zooplankton and gut microbiota. BMC genomics. 2021;22:1–18.

55. Liu F, Borg Olesen K, Reimer Borregaard A, Vollertsen J. Microplastics in urban and highway stormwater retention ponds. Science of The Total Environment. 2019;671:992–1000. 10.1016/j.scitotenv.2019.03.416

56. Liu Z, Cai M, Wu D, Yu P, Jiao Y, Jiang Q, et al. Effects of nanoplastics at predicted environmental concentration on Daphnia pulex after exposure through multiple generations. Environmental Pollution. 2020;256:113506. 10.1016/j.envpol.2019.113506Ma

57. Lobelle D, Cunliffe M. Early microbial biofilm formation on marine plastic debris. Marine pollution bulletin. 2011;62(1):197–200.

58. Love MI, Huber W, Anders S. Moderated estimation of fold change and dispersion for RNA-seq data with DESeq2. Genome Biology. 2014;15:550. 10.1186/s13059-014-0550-8

59. Macke E, Callens M, De Meester L, et al. Host-genotype dependent gut microbiota drives zooplankton tolerance to toxic cyanobacteria. Nat Commun. 2017;8:1608. 10.1038/s41467-017-01714-x

60. Macke E, Callens M, Massol F, et al. Diet and genotype of an aquatic invertebrate affect the composition of free-living microbial communities. Frontiers in Microbiology. 2020. 10.3389/fmicb.2020.00380

61. McFall A, Coughlin SA, Hardiman G, Megaw J. Strategies for biofilm optimization of plastic-degrading microorganisms and isolating biofilm formers from plastic-contaminated environments. Sustainable Microbiology. 2024. 10.1093/sumbio/qvae012

62. McMurdie PJ, Holmes S. phyloseq: An R package for reproducible interactive analysis and graphics of microbiome census data. PLOS ONE. 2013;8(4). 10.1371/journal.pone.0061217

63. Meyers N, Bouwens J, Catarino AI, De Witte B, Everaert G. Extraction of microplastics from marine seawater samples followed by Nile red staining. In: De Witte B, Power OP, Fitzgerald E, Kopke K, editors. ANDROMEDA Portfolio of Microplastics Analyses Protocols. JPI Oceans ANDROMEDA Project. 2024a. 10.13140/RG.2.2.21010.06088

64. Meyers N, De Witte B, Catarino AI, Everaert G. Automated microplastic analysis: Nile red staining and random forest modelling. In: De Witte B, Power OP, Fitzgerald E, Kopke K, editors. ANDROMEDA Portfolio of Microplastics Analyses Protocols. JPI Oceans ANDROMEDA Project. 2024c. 10.13140/RG.2.2.21010.06088

65. Morét-Ferguson S, Law KL, Proskurowski G, Murphy EK, Peacock EE, Reddy CM. The size, mass, and composition of plastic debris in the western North Atlantic Ocean. Mar Pollut Bull. 2010;60(10):1873–8. doi: 10.1016/j.marpolbul.2010.07.020. Epub 2010 Aug 14. PMID: 20709339.

66. Narmon AS, Jenisch LM, Pitet LM, Dusselier M. Ring-Opening Polymerization Strategies for Degradable Polyesters. In: Dusselier M, Lange JP, editors. Biodegradable Polymers in the Circular Plastics Economy. 2022. Available from: 10.1002/9783527827589.ch7

67. Nava V, Chandra S, Aherne J, et al. Plastic debris in lakes and reservoirs. Nature. 2023;619:317–322. 10.1038/s41586-023-06168-4

68. Oksanen J, Blanchet FG, Kindt R, Legendre P, Minchin PR, O’Hara RB, et al. vegan: Community Ecology Package. R package version 2.0-10. Available online at: http://CRAN.R-project.org/package=vegan

69. Paul MJ, Meyer JL. Streams in the urban landscape. Annual Review of Ecology and Systematics. 2001;32:333–365. DOI: 10.1146/annurev.ecolsys.32.081501.114040

70. Peerakietkhajorn S, Kato Y, Kasalicky V, Matsuura T, Watanabe H. Betaproteobacteria Limnohabitans strains increase fecundity in the crustacean *Daphnia magna*: Symbiotic relationship between major bacterioplankton and zooplankton in freshwater ecosystem. Environmental Microbiology. 2016;18(9):2366–2374.

71. Peerakietkhajorn S, Tsukada K, Kato Y, Matsuura T, Watanabe H. Symbiotic bacteria contribute to increasing the population size of a freshwater crustacean, *Daphnia magna*. Environmental Microbiology Reports. 2015;7(2):364–372. 10.1111/1758-2229.12260

72. Qi W, Nong G, Preston JF, Ben-Ami F, Ebert D. Comparative metagenomics of *Daphnia* symbionts. BMC Genom. 2009;10:21.

73. Quast C, Pruesse E, Yilmaz P, Gerken J, Schweer T, Yarza P, et al. The SILVA ribosomal RNA gene database project: improved data processing and web-based tools. Nucleic Acids Research. 2013;41(D1). 10.1093/nar/gks1219

74. Raju M, Gandhimathi R, Nidheesh PV. The cause, fate and effect of microplastics in freshwater ecosystem: Ways to overcome the challenge. Journal of Water Process Engineering. 2023;55:104199. 10.1016/j.jwpe.2023.104199

75. Rehse S, Kloas W, Zarfl C. Short-term exposure with high concentrations of pristine microplastic particles leads to immobilisation of Daphnia magna. Chemosphere. 2016 Jun;153:91–9. doi: 10.1016/j.chemosphere.2016.02.133. Epub 2016 Mar 21. PMID: 27010171.

76. Rist S, Baun A, Hartmann NB. Ingestion of micro- and nanoplastics in *Daphnia magna* – Quantification of body burdens and assessment of feeding rates and reproduction. Environmental Pollution. 2017;228:398–407. 10.1016/j.envpol.2017.05.048

77. Rodrigues MO, Abrantes N, Gonçalves FJM, Nogueira H, Marques JC, Gonçalves AMM. Spatial and temporal distribution of microplastics in water and sediments of a freshwater system (Antuã River, Portugal). Science of the total environment. 2018;633:1549–1559.

78. Rummel CD, Jahnke A, Gorokhova E, Kühnel D, Schmitt-Jansen M. Impacts of biofilm formation on the fate and potential effects of microplastic in the aquatic environment. Environmental Science & Technology Letters. 2017;4(7):258–267. 10.1021/acs.estlett.7b00164

79. Scherer C, Weber A, Stock F, Vurusic S, Egerci H, Kochleus C, et al. Comparative assessment of microplastics in water and sediment of a large European river. Science of The Total Environment. 2020;738:139866. 10.1016/j.scitotenv.2020.139866

80. Sison-Mangus MP, Mushegian AA, Ebert D. Water fleas require microbiota for survival, growth and reproduction. The ISME journal. 2015;9(1):59–67.

81. Seeley ME, Song B, Passie R, et al. Microplastics affect sedimentary microbial communities and nitrogen cycling. Nat Commun. 2020;11:2372. 10.1038/s41467-020-16235-3

82. Sruthy S, Ramasamy EV. Microplastic pollution in Vembanad Lake, Kerala, India: the first report of microplastics in lake and estuarine sediments in India. Environmental pollution. 2017;222:315-322.

83. Sullam KE, Pichon S, Schaer TM, Ebert D. The combined effect of temperature and host clonal line on the microbiota of a planktonic crustacean. Microb Ecol. 2018;76:506–517.

84. Sun Y, Duan C, Cao N, Li X, Li X, Chen Y, et al. Effects of microplastics on soil microbiome: The impacts of polymer type, shape, and concentration. Science of The Total Environment. 2022;806(Part 2):150516. DOI: 10.1016/j.scitotenv.2021.150516

85. Szabó I, Al-Omari J, Szerdahelyi GS, et al. In Situ Investigation of Plastic-Associated Bacterial Communities in a Freshwater Lake of Hungary. Water Air Soil Pollut. 2021;232:493. 10.1007/s11270-021-05445-0

86. Tackmann J, Rodrigues JFM, von Mering C. Rapid inference of direct interactions in large-scale ecological networks from heterogeneous microbial sequencing data. Cell Systems. 2019;9(3):286–296.e8. 10.1016/j.cels.2019.08.002

87. Trotter B, Wilde MV, Brehm J, Dafni E, Aliu A, Arnold GJ, et al. Long-term exposure of *Daphnia magna* to polystyrene microplastic (PS-MP) leads to alterations of the proteome, morphology and life-history. Sci Total Environ. 2021 Nov 15;795:148822. doi: 10.1016/j.scitotenv.2021.148822. Epub 2021 Jul 3. PMID: 34328913

88. Tian L, van Putten RJ, Gruter GJM. Plastic Pollution. The Role of (Bio)Degradable Plastics and Other Solutions. In: Dusselier M, Lange JP, editors. Biodegradable Polymers in the Circular Plastics Economy. 2022. Available from: 10.1002/9783527827589.ch3

89. Vercauteren M, Semmouri I, van Acker E, Pequeur E, van Esch L, Uljee I, et al. Kernrapport-microplasticonderzoek-finaalUGent_TW. 2021.

90. Vlaanderen EJ, Ghaly TM, Moore LR, Focardi A, Paulsen IT, Tetu SG. Plastic leachate exposure drives antibiotic resistance and virulence in marine bacterial communities. Environmental Pollution. 2023;327:121558. 10.1016/j.envpol.2023.121558

91. Wang M, Wang WX. Selective ingestion and response by *Daphnia magna* to environmental challenges of microplastics. Journal of Hazardous Materials. 2023;458:131864. 10.1016/j.jhazmat.2023.131864

92. Wood DE, Lu J, Langmead B. Improved metagenomic analysis with Kraken 2. Genome Biology. 2019;20:257. 10.1186/s13059-019-1891-0

93. Xiong X, et al. Sources and distribution of microplastics in China’s largest inland lake – qinghai lake. Environmental Pollution. 2018;235:899–906. doi:10.1016/j.envpol.2017.12.081.

94. Yuan W, et al. Microplastic abundance, distribution and composition in water, sediments, and wild fish from Poyang Lake, China. Ecotoxicology and Environmental Safety. 2019;170:180–187. doi:10.1016/j.ecoenv.2018.11.126.

95. Zadjelovic V, Wright RJ, Borsetto C, et al. Microbial hitchhikers harbouring antimicrobial-resistance genes in the riverine plastisphere. Microbiome. 2023;11:225. 10.1186/s40168-023-01662-3

96. Zettler ER, Mincer TJ, Amaral-Zettler LA. Life in the ‘ plastisphere ’: microbial communities on plastic marine debris. Environ Sci Technol. 2013;47:7137–7146. 10.1021/es401288x

97. Zhang K, Su J, Xiong X, Wu X, Wu C, Liu J. Microplastic pollution of lakeshore sediments from remote lakes in Tibet plateau, China. Environmental pollution. 2016;219:450–455.

98. Zhao K, Li C, Li F. Research progress on the origin, fate, impacts and harm of microplastics and antibiotic resistance genes in wastewater treatment plants. Sci Rep. 2024;14:9719. 10.1038/s41598-024-60458-z

99. Zhao K, Li C, Li F. Research progress on the origin, fate, impacts and harm of microplastics and antibiotic resistance genes in wastewater treatment plants. Sci Rep. 2024;14:9719. 10.1038/s41598-024-60458-z

100. Zhu D, Ma J, Li G, et al. Soil plastispheres as hotspots of antibiotic resistance genes and potential pathogens. ISME J. 2022;16:521–532. 10.1038/s41396-021-01103-9

101. Zimmermann L, Göttlich S, Oehlmann J, Wagner M, Völker C. What are the drivers of microplastic toxicity? Comparing the toxicity of plastic chemicals and particles to *Daphnia magna*. Environmental Pollution. 2020;267:115392. 10.1016/j.envpol.2020.115392

