## Supplementary materials and methods for "Effects of microplastics on Daphnia-associated microbiomes in situ and in vitro"

**Supplementary materials & methods**

1. **Characterisation and quantification of MP from sediment and water samples**

The methods are based on the protocol developed by Meyers et al., (2024).

Briefly, water samples were filtered over a metal 20 µm mesh size sieve. The residue was rinsed off the filter with demineralised water into a glass beaker and the organic matter was digested with 100 ml of H_2_O_2_ (33%) at 60°C and 150 rpm for 48h. The digested solution was transferred back onto the sieve and rinsed thoroughly with 50mL demineralised water. If the original water sample was mixed with sediment, three density separation steps with saturated sodium iodide solution (NaI – 1.8 g/cm³) were necessary to extract all microplastic particles (Frais *et al*. 2018).

To isolate MP particles from sediment samples, the samples were first dried in an oven at 50°C for 72h. From 50 g of dried sediment, all MP particles were extracted by three density separation steps using a saturated sodium iodide solution (NaI – 1.8 g/cm³, Frais *et al*. 2018). Similar to water samples, the organic matter was removed by adding H_2_O_2_ (33%) followed by 1-week incubation at a room temperature. The residue was filtered over 20 µm mesh sieve and rinsed as above.

After the organic matter digestion, the number of MP was counted for each sample. Here, the demineralised water containing MP was filtered over a 10 µm mesh PTFE membrane (Omnipore) using a manifold set with a vacuum pump. Thereafter, the PTFE filter was stained with ml Nile Red (10 µg/ml acetone). After 15 minutes, the filter was washed with demineralized water and left for 24 h to dry. A fluorescence microscope (Leica DM 1000, LAS Core software) was used to obtain images of Nile Red-dyed particles for each sample. A two-step semi-automated decision tree classification model as described by Meyers et al., (2022), was used to both distinguish plastic from non-plastic particles (Plastic Detection Model, PDM) and for plastic polymer identification (Polymer Identification Model, PIM).

Positive control samples were prepared by spiking MP particles in the matrix (composed of milliQ water and purified seasand) with heterogeneously shaped polyethylene (PE), polyethylene terephthalate (PET), polypropylene (PP), polystyrene (PS), and polyvinyl chloride (PVC) fragments provided by Carat PLC (Westerlo, Belgium) at following sizes: 500-1000 μm and 100-300 μm.

2. **Amplification of V4 region 16rRNA gene for Illumina Miseq sequencing**

Initially, the full-length 16S rRNA gene was amplified using forward EUB8F and reverse 1492R primers (**Table 1**) with 10 ng of template DNA and high-fidelity SuperFi polymerase (Life Technologies) across 30 cycles of denaturation at 98°C for 10 seconds, annealing at 50°C for 45 seconds, and extension at 72°C for 30 seconds. The PCR products were purified using the CleanPCR kit (CleanNA), according to the manufacturer's instructions, and DNA concentrations were determined with a Nanodrop spectrophotometer. A second amplification was then conducted on 10 ng of the purified PCR product using forward 515F and reverse 806R primer pair (**Table 1**) primer pair for 30 cycles, under the same thermal cycling conditions as before (Kozich et al., 2013). The second step PCR primers incorporated both an Illumina adapter sequence and an 8-nucleotide barcode at the 5' end for multiplex sequencing. The last PCR amplifications reactions were

conducted in triplicate for each sample, pooled, and the correct size of the amplicon verified on a 1.5% agarose gel. The amplicons were purified using the CleanPCR kit (CleanNA) and sequenced on the Illumina MiSeq platform using a v2 PE500 kit producing 250-nt paired-end reads. To prepare an equimolar library for sequencing, amplicon concentrations were normalised using a SequalPrep Normalisation Plate (Applied Biosystems) and pooled accordingly.

**Table 1. Primers for external and internal PCR**

| Primer name | Primer sequence |
| --- | --- |
| EUB8F | 5’ – AGA GTT TGA TCM TGG CTC AG – 3’ |
| 1492R | 5’ – GGT TAC CTT GTT ACG ACT T – 3’ |
| 515F | 5’ – AATGATACGGCGACCACCGAGATCTACAC-atcgtacgTATGGTAATTGTGTGCCAGCMGCCGCGGTAA – 3’ |
| 806R | 5’ – CAAGCAGAAGACGGCATACGAGAT-actatgtc AGTCAGTCAGCCGGACTACNVGGGTWTCTAAT – 3’ |

3. **Making microfibers using cryotome cutting machine**

The fibres were embedded in a freezing agent (Epredia™ Neg-50™ Frozen Section Medium, Fisher Scientific #12698086) and solidified at -80°C. Thereafter, the blocks were cut into 50 µm pieces aligned perpendicularly to the cryotome blade. The resulting microfibers were collected on tin foil, submerged in MQ water, heated at 60°C for 1h in the oven and subsequently collected via vacuum filtration. The fibres were examined under a light microscope (Olympus BX51) to record shape and diameter.

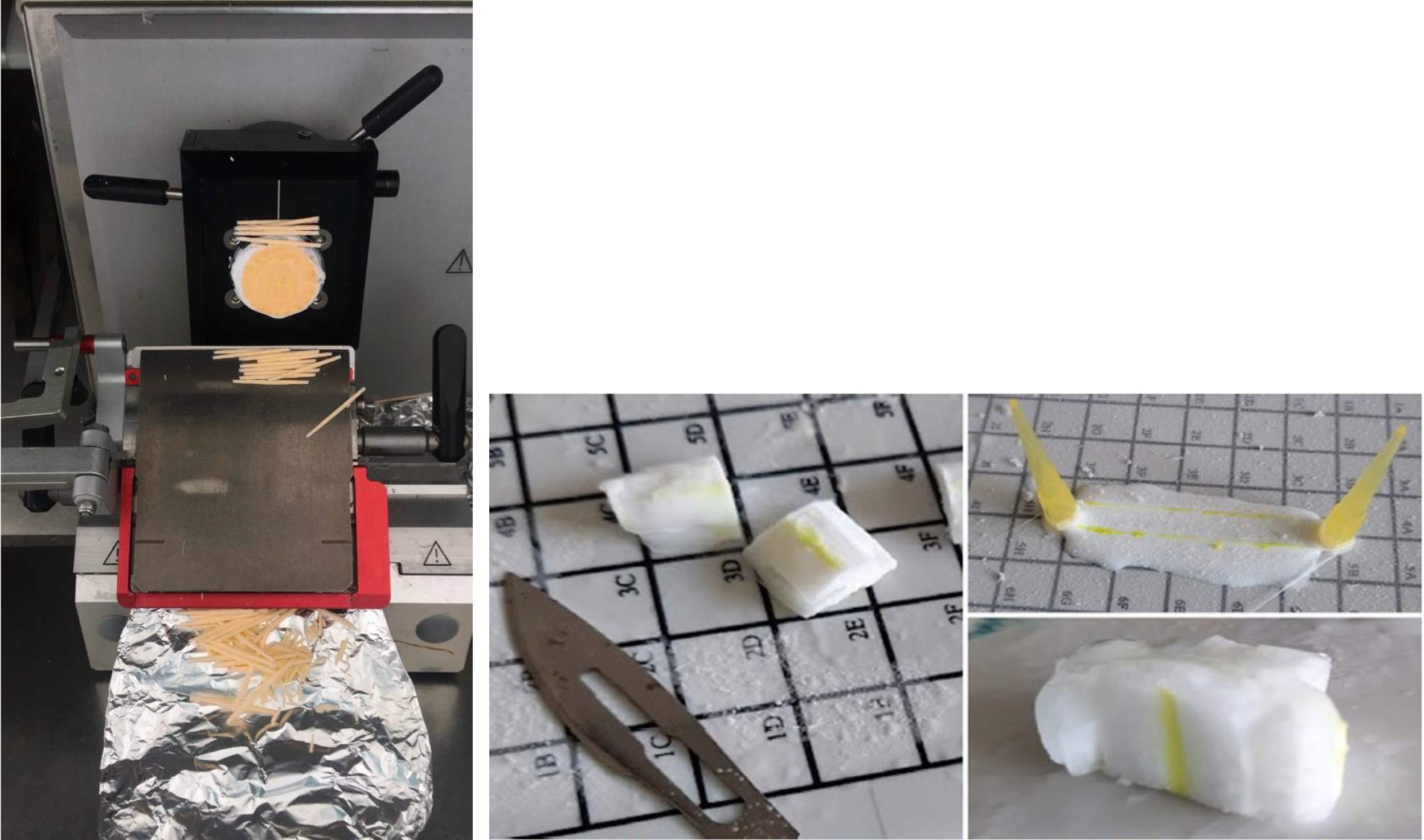

**Figure 1.** The fiber was wrapped around pipette tip and covered with a freezing agent (Right). All pieces were then cut and glued together (Right). A block was formed with all the pieces (Right), the fibers all point in the same direction to be placed perpendicular to the blade of the cryotome (Left).

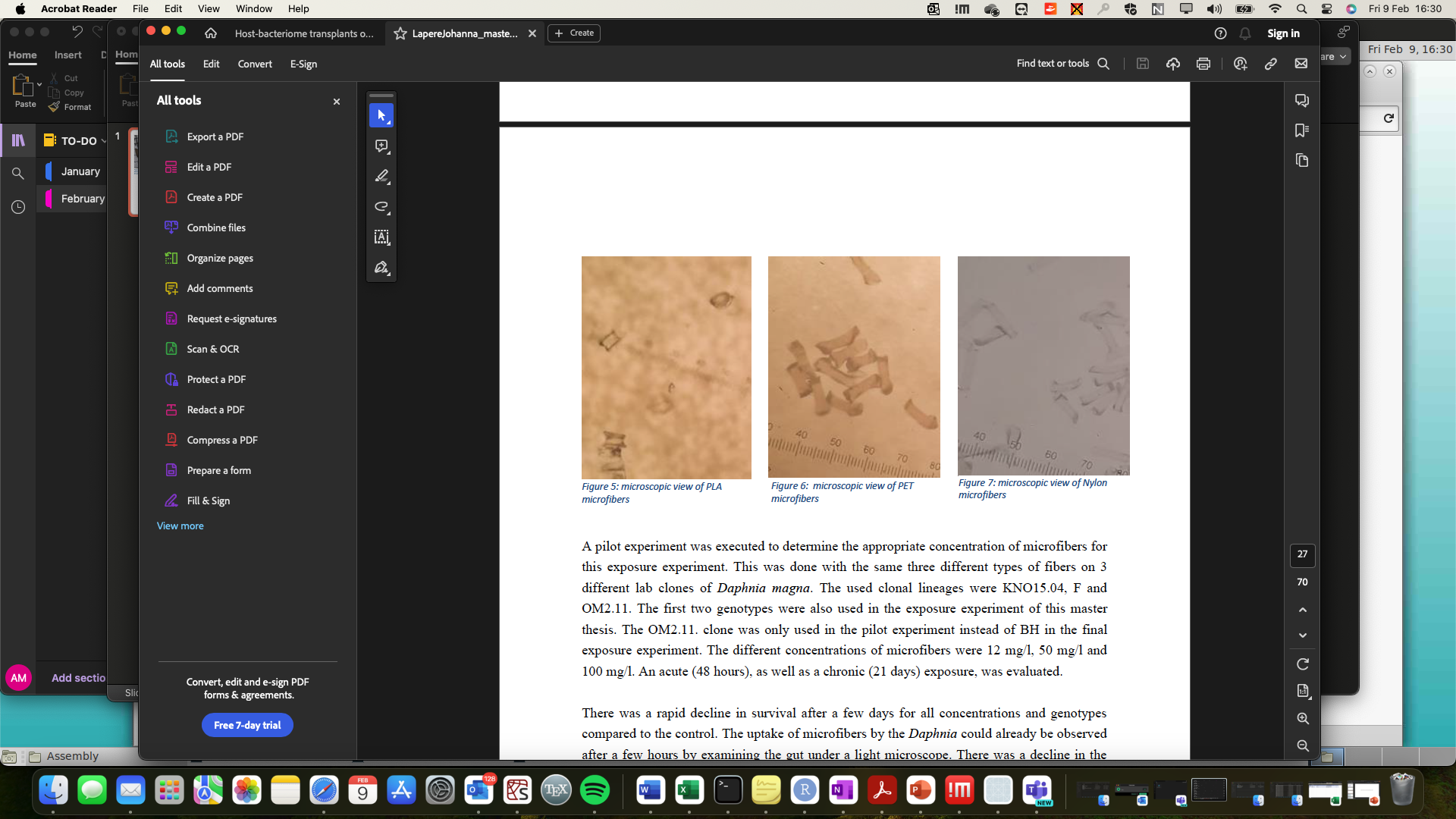

Nylon

PET

PLA

**Figure 2**. Microscopic photographs of MP generated for the study.

4. **Dead/Alive plots for PET exposure at different concentrations (mg/L) and different *Daphnia magna* clones (KNO 15.05, OM 2.11, Line F).**

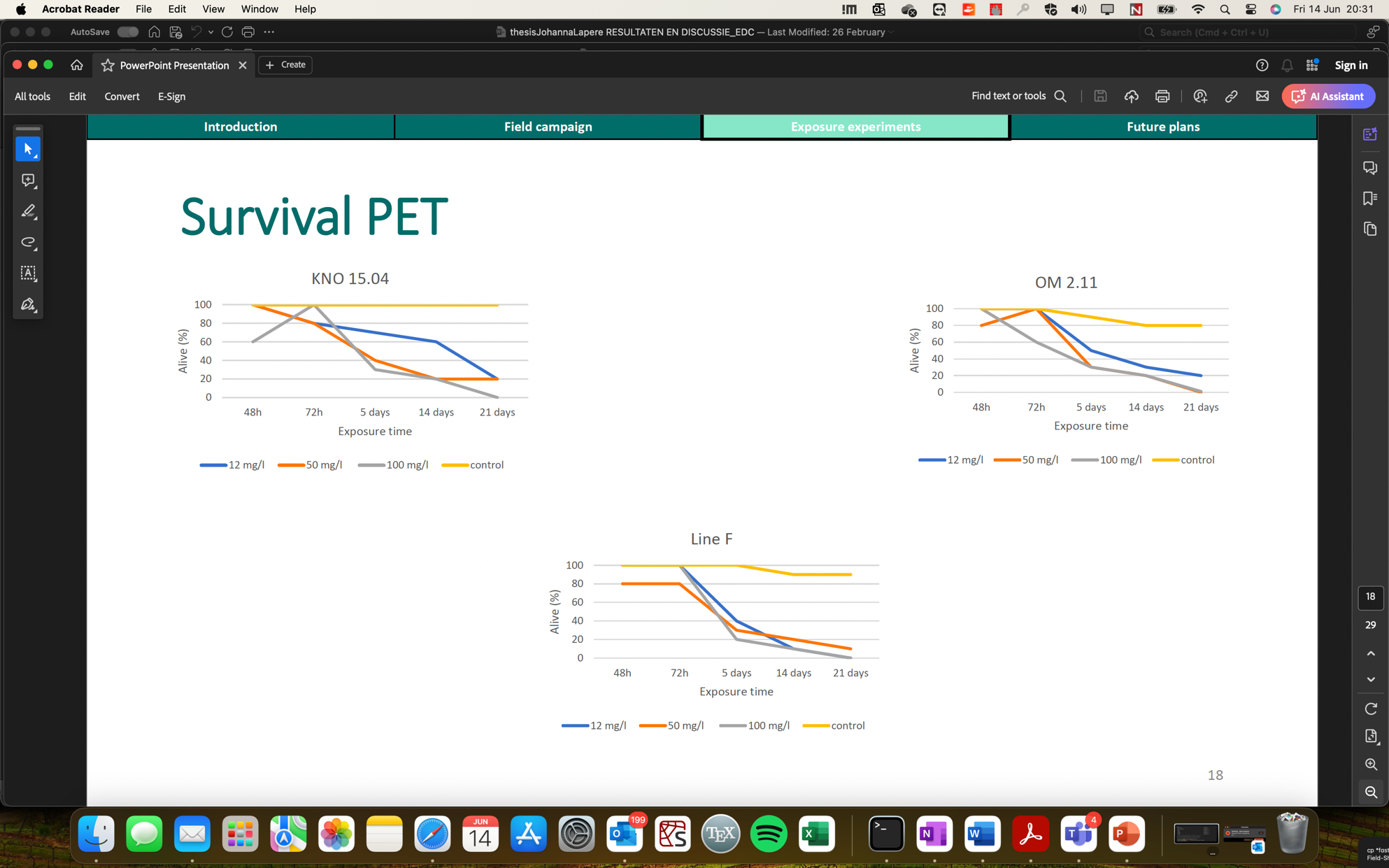

5. **Daphnia magna (KNO 15.04) body size after 14 and 21 days exposure to varying concentrations of PLA and Nylon 6.**

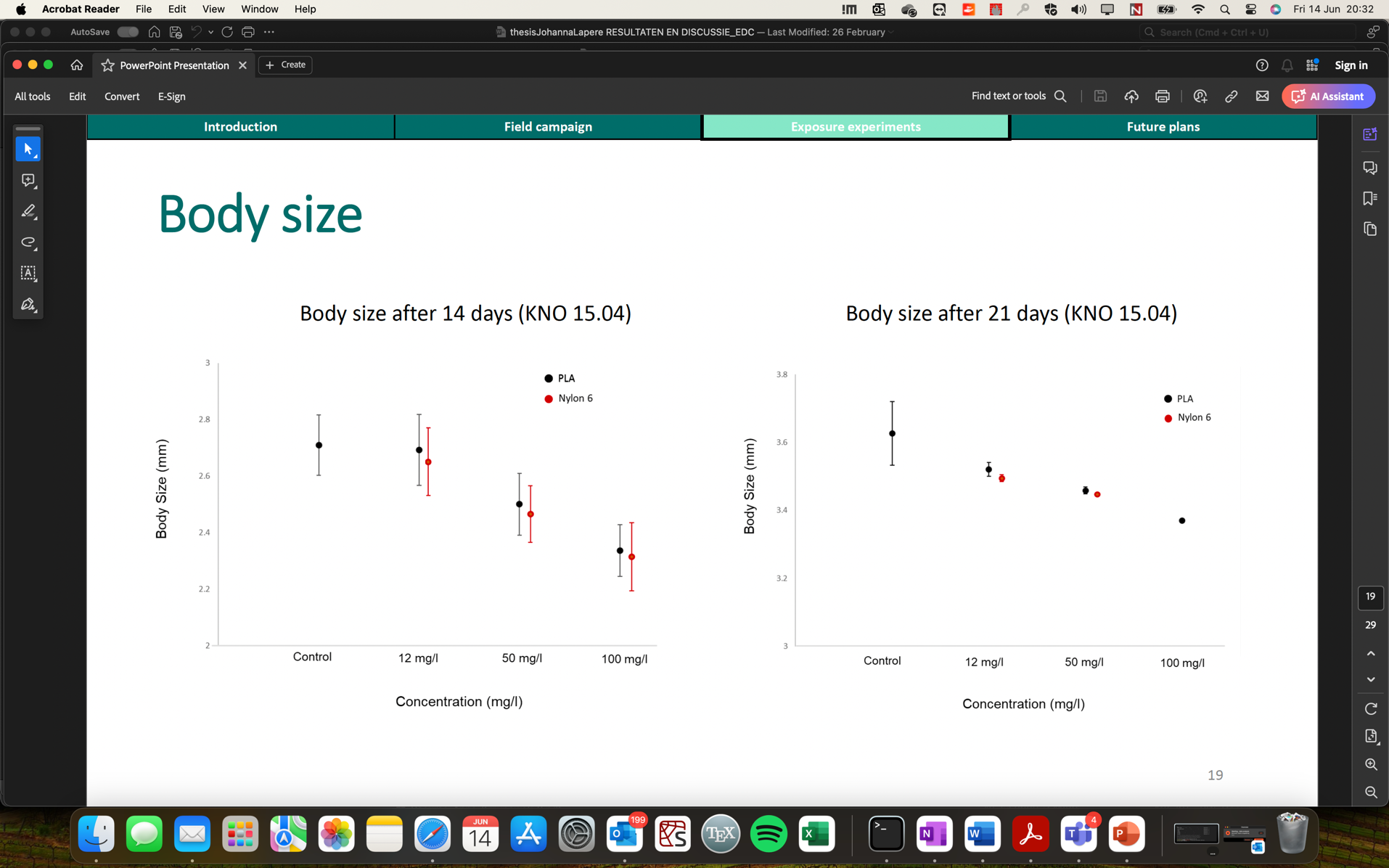

6. **Measured parameters of Blauwe Poort (BP) and De-Gavers (DG) inoculums before the MP exposure experiment**

| **Pond** | **°C** | **dissolved oxygen (mg/l)** | **Liquid dissolved oxygen (%)** | **pH** | **Conductivity (µs/cm)** | **Redox (mV)** |
| --- | --- | --- | --- | --- | --- | --- |
| **DG1** | 11.7 | 9.39 | 85.5 | 8.05 | 752 | 174.3 |
| **DG2** | 11.6 | 9.39 | 85.2 | 8.05 | 752 | 174.2 |
| **DG3** | 11.6 | 9.36 | 84.8 | 8.05 | 751 | 174.6 |
| **BP1** | 9.5 | 1.69 | 14.6 | 7.84 | 840 | -12.2 |
| **BP2** | 9.4 | 1.59 | 13.7 | 7.84 | 846 | -9.8 |
| **BP3** | 9.3 | 1.53 | 13.2 | 7.84 | 845 | -8 |

7. ***Daphnia magna* MP exposure jar set-up**

| \| Water medium \| \| Sterilized water \| Sterilized water; \| Microfiber type \| \| No microfibers \| No microfibers; \| Genotype \| \| No *Daphnia* \| No *Daphnia* | | **PET** | **NYLON** | **PLA** | **NO MF** |
| --- | --- | --- | --- | --- | --- |
| **Blauwe P** | F | Blauwe P + F + PET | Blauwe P + F + NYLON | Blauwe P + F + PLA | Blauwe P + F |
|  | KNO 15.04 | Blauwe P + KNO 15.04 + PET | Blauwe P + KNO 15.04 + NYLON | Blauwe P + KNO 15.04 + PLA | Blauwe P + KNO 15.04 |
|  | BH | Blauwe P +  BH + PET | Blauwe P + BH + NYLON | Blauwe P + BH + PLA | Blauwe P + BH |
|  | NO *DAPHNIA* | Blauwe P + PET | Blauwe P + NYLON | Blauwe P + PLA |  |
| **De Gavers** | F | De Gavers + F + PET | De Gavers + F + NYLON | De Gavers + F + PLA | De Gavers + F |
|  | KNO 15.04 | De Gavers + KNO 15.04 + PET | De Gavers + KNO 15.04 + NYLON | De Gavers + KNO 15.04 + PLA | De Gavers + KNO 15.04 |
|  | BH | De Gavers +  BH + PET | De Gavers +  BH + NYLON | De Gavers +  BH + PLA | De Gavers +  BH |
|  | NO *DAPHNIA* | De Gavers + PET | De Gavers + NYLON | De Gavers + PLA |  |
| **Sterilised H_2_O** | NO *DAPHNIA* | Sterilised H_2_O + PET | Sterilised H_2_O + NYLON | Sterilised H_2_O + PLA |  |

8. **Stereoscopic photo showing a randomly picked *Daphnia magna* sample with fibres embedded in the gut tissues.**

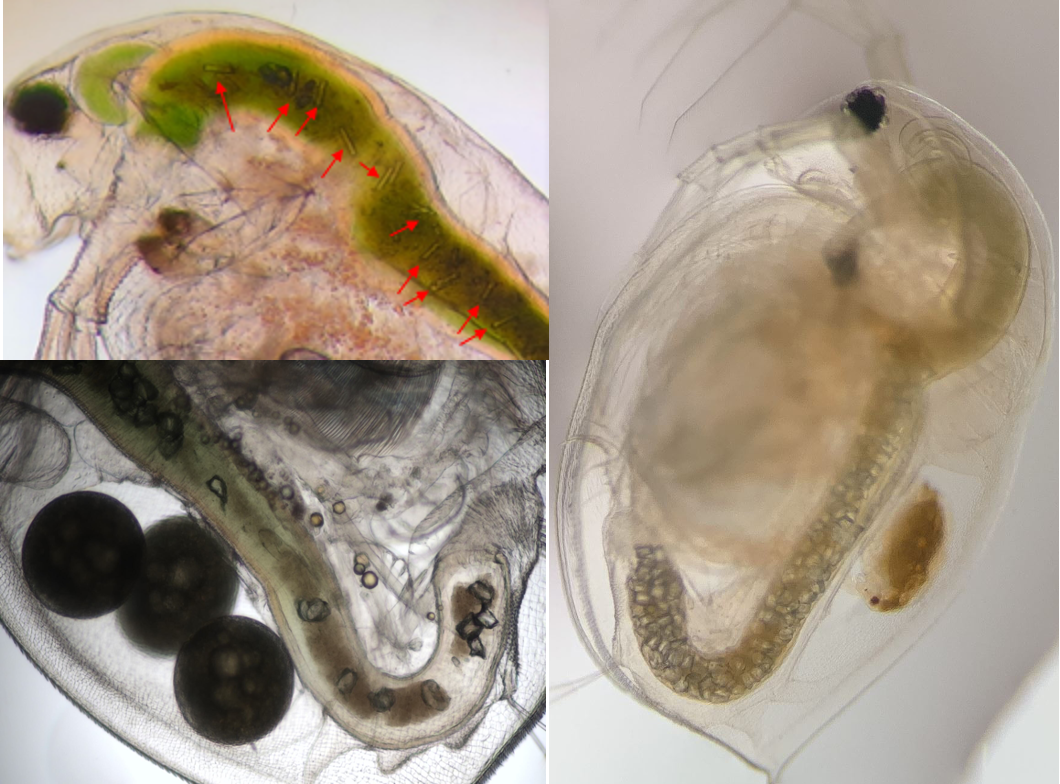

References:

Frais, J., Pagter, E., Nash, R., O'Connor, I., Carretero, O., Filgueiras, A., Viñas, L., Gago, J., Antunes, J., Bessa, F., Sobral, P., Goruppi, A., Tirelli, V., Pedrotti, M. L., Suaria, G., Aliani, S., Lopes, C., Raimundo, J., Caetano, M., & Gerdts, G. (2018). Standardised protocol for monitoring microplastics in sediments. https://doi.org/10.13140/RG.2.2.36256.89601/1

Kozich, J. J., Westcott, S. L., Baxter, N. T., Highlander, S. K., & Schloss, P. D. (2013). Development of a dual-index sequencing strategy and curation pipeline for analyzing amplicon sequence data on the MiSeq Illumina sequencing platform. Applied and Environmental Microbiology, 79(17), 5112–5120. https://doi.org/10.1128/AEM.01043-13

Meyers, N., De Witte, B., Catarino, A. I., & Everaert, G. (2024). Extraction of microplastics from marine sediment samples followed by Nile red staining. In B. De Witte, O.-P. Power, E. Fitzgerald, & K. Kopke (Eds.), ANDROMEDA Portfolio of Microplastics Analyses Protocols. ANDROMEDA Deliverable 5.5. JPI Oceans ANDROMEDA Project. <https://doi.org/10.13140/RG.2.2.21010.06088>

Meyers N, Catarino AI, Declercq AM, Brenan A, Devriese L, Vandegehuchte M, et al. Microplastic detection and identification by Nile red staining: Towards a semi-automated, cost- and time-effective technique. Sci Total Environ. 2022;823:153441. doi:10.1016/j.scitotenv.2022.153441.
