## Supplementary Tables and Figures for "Effects of microplastics on Daphnia-associated microbiomes in situ and in vitro"

**Supplementary Figures and Tables**

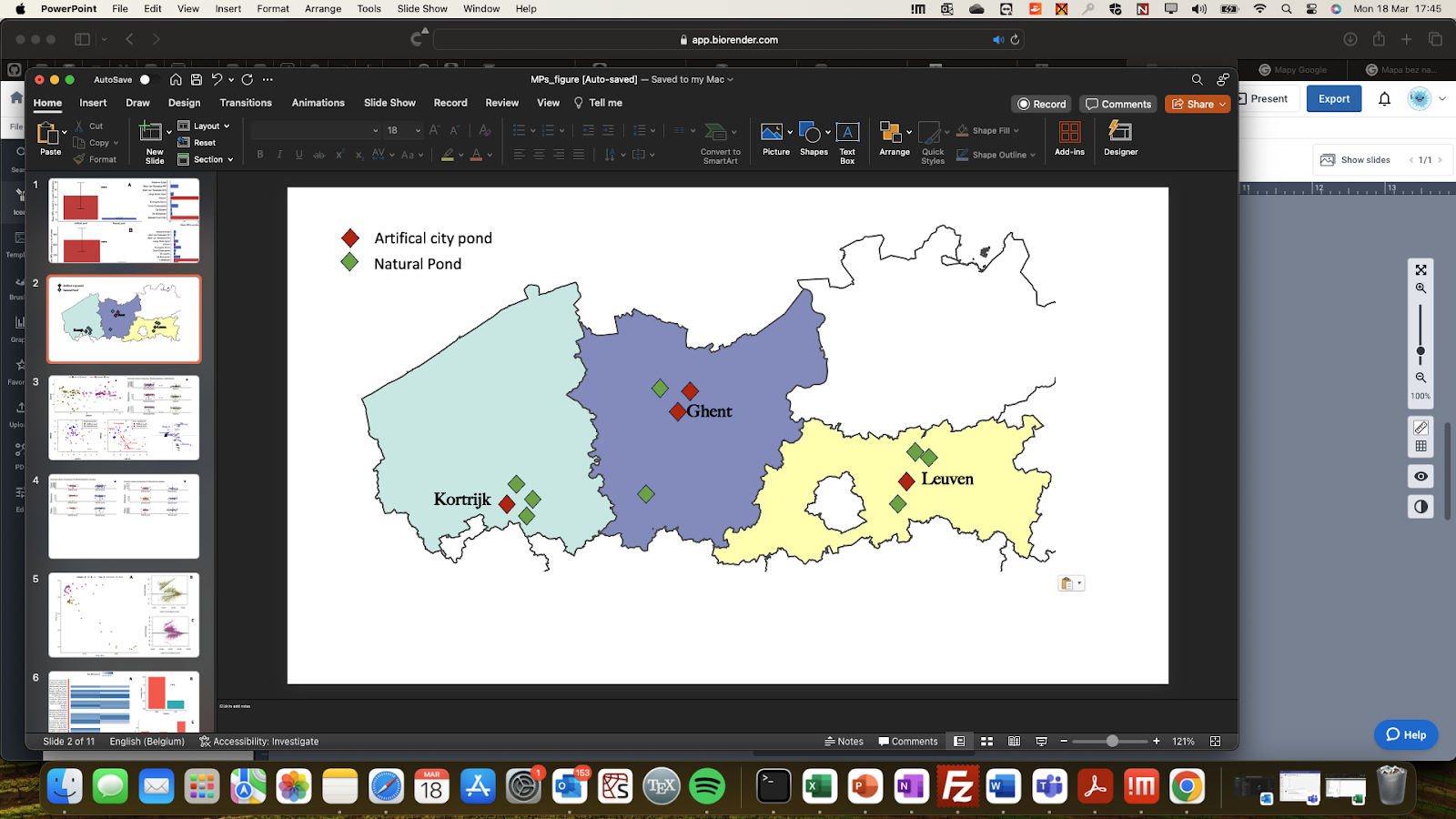

**Supplementary figure 1. Map showing geographical locations of sampled ponds in Flanders, Belgium during the sampling campaign of 2021.** City ponds are marked in dark-red and natural ponds are marked in green. Major cities considered in the study are labelled.

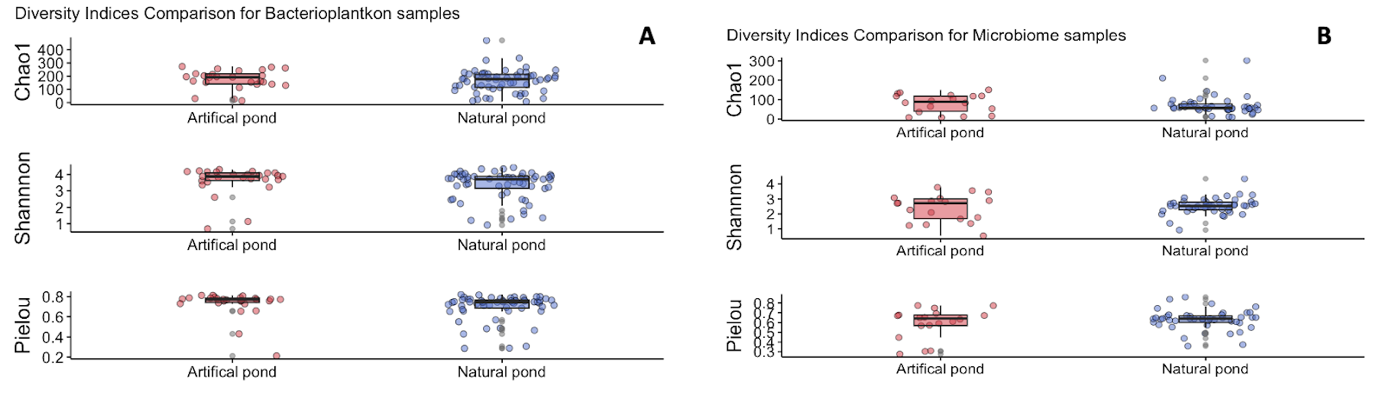

**Supplementary figure 2. Diversity indices for Bacterioplankton (A) and Microbiome field samples (B).**

**Supplementary table 1.**

*Taxa-MP associations inferred from a microbial network for Daphnia magna host-microbiomes exposed in-vitro to MP (Nylon, PET, PLA). Colour corresponds to the sign of association, positive-red and negative-blue.*

| Plastic | Taxa | Taxa origin | Weight | 16S-sequence |
| --- | --- | --- | --- | --- |
| PET | Prosthecobacter | Daphnia  Core + Incolulum | 0.33693376 | AATCACTGGGCGTAAAGGGTGCGTAGGTGGCTTGGTAAGTCAGATGTGAAAGCCCGGGGCTCAACCTCGGAATTGCATCCGATACTGCTAGGCTAGAGTACTGGAGGGGTGACTAGAATTCTCGGTGTAGCAGTGAAATGCGTAGATATCGAGAGGAATACCAAAGGCGTAGGCAGGTCACTGGACAGTTACTGACACTGAGGCACGAAGGCCAG |
| Nylon | Planctomycetales | Inoculum | 0.38520375 | AATCACTGGGCTTAAAGGGTGCGTAGGCGGTCTTTTAAGTAGGGTGTGAAAGCCTCTGGCTCAACCAGAGAACTGCGCCCTAAACTAGAAGGCTTGAGTGAGGTAGGGGTGTGTGGAACTTCCAGTGGAGCGGTGAAATGTGTTGATATTGGAAGGAACGCCGGTGGCGAAAGCGACACACTGGACCTTGTCTGACGCTGAGGCACGAAAGCCAG |
| Nylon | Gemmobacter | Inoculum | 0.24894698 | AATTACTGGGCGTAAAGCGCACGTAGGCGGATCAGAAAGTCAGAGGTGAAATCCCAGGGCTCAACCTTGGAACTGCCTTTGAAACTCCTGGTCTTGAGGTCGAGAGAGGTGAGTGGAATTCCGAGTGTAGAGGTGAAATTCGTAGATATTCGGAGGAACACCAGTGGCGAAGGCGGCTCACTGGCTCGATACTGACGCTGAGGTGCGAAAGCGTG |
